# Reference-guided genome assembly at scale using ultra-low-coverage high-fidelity long-reads with HiFiCCL

**DOI:** 10.1101/2025.04.20.649739

**Authors:** Zhongjun Jiang, Weihua Pan, Runtian Gao, Heng Hu, Wentao Gao, Murong Zhou, Yu-Hang Yin, Zhipeng Qian, Shuilin Jin, Guohua Wang

**Affiliations:** College of Life Science, Northeast Forestry University, Harbin 150000, China; College of Computer and Control Engineering, Northeast Forestry University, Harbin 150000, China; Shenzhen Branch, Guangdong Laboratory of Lingnan Modern Agriculture, Genome Analysis Laboratory of the Ministry of Agriculture and Rural Affairs, Agricultural Genomics Institute at Shenzhen, Chinese Academy of Agricultural Sciences, Shenzhen 518120, China; School of Mathematics, Harbin Institute of Technology, Harbin 150001, China; School of Computer Science and Technology, Harbin Institute of Technology, Harbin 150001, China

**Keywords:** reference-guided assembly, ultra-low coverage, long high-fidelity reads, chromosome-by-chromosome, population genomics

## Abstract

Population genomics using short-read resequencing captures single nucleotide polymorphisms and small insertions and deletions but struggles with structural variants (SVs), leading to a loss of heritability in genome-wide association studies. In recent years, long-read sequencing has improved pangenome construction for key eukaryotic species, addressing this issue to some extent. Sufficient-coverage high-fidelity (HiFi) data for population genomics is often prohibitively expensive, limiting its use in large-scale populations and broader eukaryotic species and creating an urgent need for robust ultra-low coverage assemblies. However, current assemblers underperform in such conditions. To address this, we propose HiFiCCL, the first assembly framework specifically designed for ultra-low-coverage high-fidelity reads, using a reference-guided, chromosome-by-chromosome assembly approach. We demonstrate that HiFiCCL improves ultra-low-coverage assembly performance of existing assemblers and outperforms the state-of-the-art assemblers on human and plant datasets. Tested on 45 human datasets (∼5x coverage), HiFiCCL combined with hifiasm reduces the length of misassembled contigs relative to hifiasm by an average of 21.19% and up to 38.58%. These improved assemblies enhance germline structural variant detection, reduce chromosome-level mis-scaffolding, enable more accurate pangenome graph construction, and improve the detection of rare and somatic structural variants based on the pangenome graph under ultra-low-coverage conditions.

## Introduction

Population genomics is crucial for understanding genetic variation within and between populations, shedding light on evolutionary processes, adaptation, and the genetic basis of traits. It has broad applications in areas such as human health, agriculture, and disease management (*1–3*). Due to the limitations of read length, the population genomes based on short-read resequencing data primarily capture information on single nucleotide polymorphisms (SNPs) and small insertions and deletions (indels) but struggle to accurately detect structural variants (SVs), leading to a loss of heritability in genome-wide-association studies (GWAS). To overcome this limitation, recent efforts have led to the construction of pangenomes for a large number of key eukaryotic species (e.g., human (*4, 5*), tomato (*6*), rice (*7*)), featuring high-quality, chromosome-level genomes assembled from high-coverage long reads. Studies have shown that with accurate and complete SV detection, pangenomes can significantly improve heritability compared to resequencing-based populations (*8–10*). However, despite a decrease in cost over the years, long-read sequencing remains significantly more expensive than short-read sequencing, limiting its application in building large-scale populations (thousands of individuals or more) and in broader eukaryotic species. Therefore, there is an urgent need for methodological innovations to achieve population-scale genome assemblies of high quality at a cost comparable to resequencing.

One possible approach to achieve this goal is to assemble ultra-low coverage (5x-10x) long reads, which are cost-effective compared to standard short-read resequencing (30x-60x), while still providing genomic sequences of sufficient accuracy, contiguity, and completeness for reliable variation detection. Based on the type of data used, existing long-read assembly algorithms can be classified into: 1) algorithms based on noisy reads (ONT or PacBio CLR), 2) algorithms based on high-fidelity (PacBio HiFi) reads, and 3) algorithms based on hybrid data (both noisy reads and HiFi reads) (*11*). The noisy-read-based assemblers (e.g. Canu (*12*), Flye (*13*), wtdbg (*14*), Nextdenovo (*15*), MECAT (*16*)) focus on addressing sequencing errors in reads to minimize their impact on the accuracy and continuity of the assembly. The HiFi-based assemblers (e.g. hifiasm (*17, 18*), HiFlye (*13*), LJA (*19*)) focus on leveraging the dual advantages of HiFi reads in accuracy and length to achieve assembly goals, such as chromosome haplotyping and the assembly of complex regions, which were previously unattainable. The hybrid assemblers (e.g. hifiasm (UL) (*20*), Verkko (*21*)) integrate the ONT ultra-long reads on the HiFi-based assembly graph to solve the ultra-complex genomic regions and can even generate chromosome-level contigs in some situations. However, nearly all these assembly methods are developed for long reads with high coverage, and our experimental results demonstrate that they struggle to achieve high-quality assembly with ultra-low coverage sequencing data, resulting in high assembly errors or incomplete assembled genomes.

A potential solution is to use a reference genome to alleviate the challenges of assembling ultra-low-coverage data. After all, when attempting to construct a population genome for a species, there is typically at least one high-quality reference genome available. Furthermore, genomics has entered the telomere-to-telomere (T2T) era, with an increasing number of high-quality reference genomes being released for various species (*11, 22–26*). This makes such as approach applicable to a much broader range of scenarios. In previous related studies, TRFill (*27*) effectively addressed the haplotype-resolved assembly of long tandem repeats by using a high-quality reference genome to bin reads from different complex genomic regions. GALA optionally utilizes a reference genome to assist multiple preliminary assemblies in guiding reads to cluster by chromosomes, enabling chromosome-by-chromosome assembly. Notably, these methods, collectively referred to as reference-guided assembly, differ from traditional reference-based assembly methods. Instead of directly assembling reads based on their alignment positions on the reference genome, they only use reference to classify the reads of different genomic regions and then perform de novo assembly. This approach not only leverages the reference genome to reduce the difficulty of assembly but also enables accurate assembly of genomic regions with large structural variations. However, despite these studies demonstrating the effectiveness of reference-guided assembly on high coverage long-reads data, their performance remains suboptimal under ultra-low-coverage conditions.

In this paper, we propose HiFiCCL, the first assembly framework specifically designed for ultra-low-coverage HiFi reads, using an exclusively reference-guided, chromosome-by-chromosome assembly approach. We validate the effectiveness of HiFiCCL across various applications, including contig-level assembly, chromosome-level scaffolding, assembly-based SVs detection, pangenome graph construction, and pangenome-graph-based rare germline and somatic SVs detection on a large number of ultra-low-coverage HiFi datasets.

## Results

### Overview of the HiFiCCL framework

Inspired by GALA, HiFiCCL designs a novel exclusively referenced-guided chromosome-by-chromosome assembly strategy, where whole-genome reads are partitioned by chromosome using the high-quality reference genome and then assembled individually. This approach can prevent interference between sequences from similar regions on different chromosomes, thereby reducing assembly difficulty and improving the quality of the assembly. The key challenge of chromosome-by-chromosome assembly lies in accurately clustering reads by chromosome. To achieve this, GALA combines multiple sets of preliminary assemblies to guide the clustering process and can optionally incorporate a reference genome as an auxiliary aid. However, in the case of ultra-low sequencing coverage, the quality of preliminary assemblies drops dramatically, with high assembly errors and incomplete genome sizes, making it difficult to accurately cluster the reads. Therefore, HiFiCCL relies solely on a high-quality reference genome to guide the read clustering process (Fig.1 and Fig. S1). Specifically, each read is aligned to the reference genome and assigned to all potential chromosome categories. However, due to sequence differences between the reference genome and the genome to be assembled, alignment errors, and highly similar regions between chromosomes, a certain proportion of reads may remain unclassified or misclassified. Therefore, HiFiCCL uses a series of innovative strategies to improve the accuracy and completeness of clustering.

**Fig. 1.**
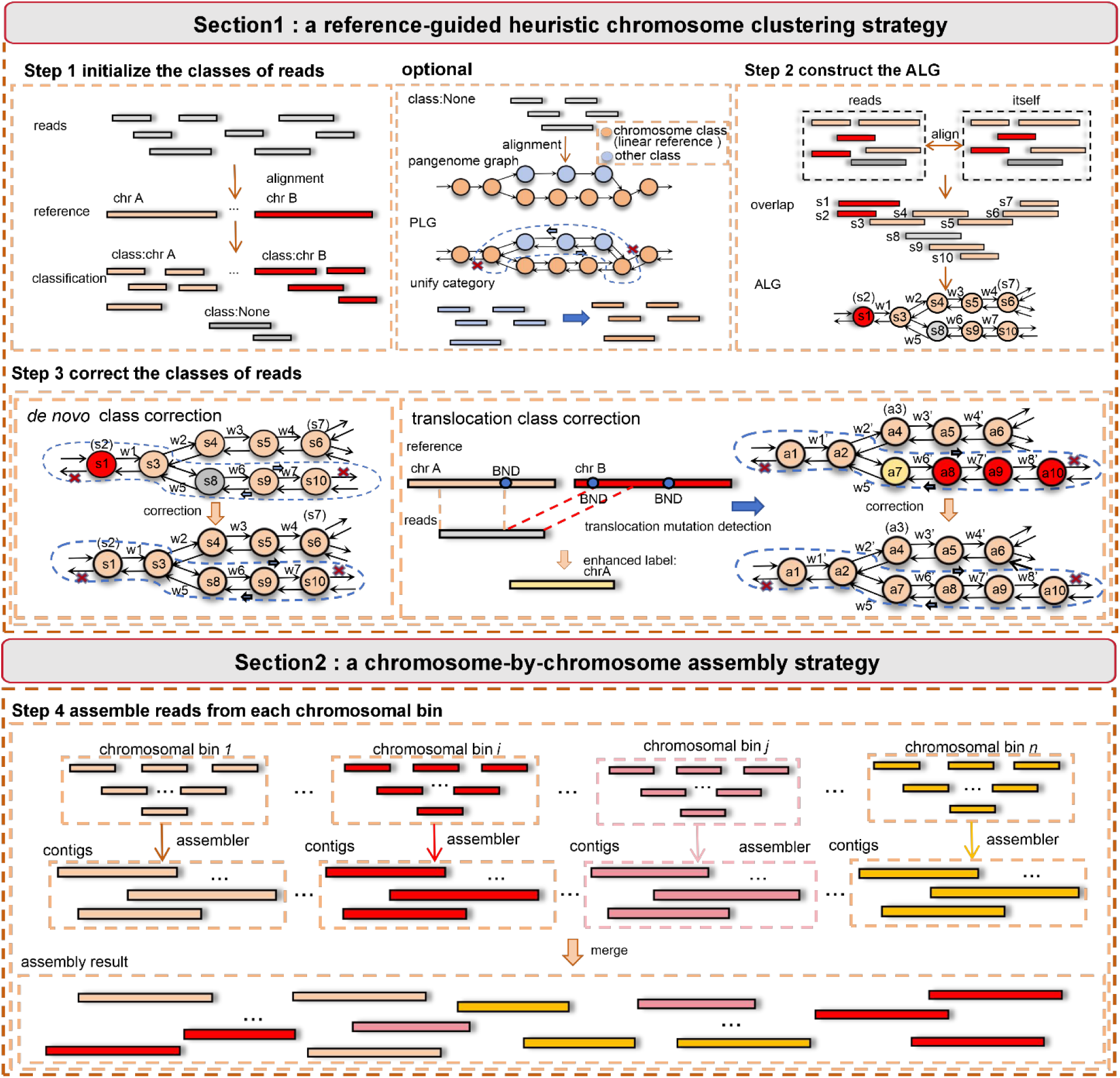
The whole pipeline of HiFiCCL. In the depicted schema, gray represents reads considered as belonging to the “None” class due to a lack of chromosomal class information, whereas other colors represent distinct chromosomal classes. HiFiCCL is structured into two sections. Section 1 encompasses three steps: initially, reads are aligned with the linear reference genome, and chromosomal classes for reads are initialized based on this alignment information. An optional mode of HiFiCCL involves aligning reads labeled as “None” to the pangenome graph. These reads are then assigned chromosomal classes by searching through the constructed Pangenome Label Graph (PLG) (see “Materials and Methods” section). The second step involves inter-read alignments from which the proposed the Alignment Label Graph (ALG) (see “Materials and Methods” section) is constructed. The third step utilizes the ALG generated from the preceding steps to correct class errors and complete the chromosome clustering process. The search begins by randomly selecting a node (e.g., node ‘s9’) and proceeds bidirectionally using a greedy approach to select the node with the highest overlap degree, until a significant drop in over-lap degree is encountered (see “Materials and Methods”). This identifies a path that serves as evidence for do novo class correction. For translocation mutations, class correction incorporates alignment information and a translocation detection algorithm to identify breakpoints. Reads containing break-points information are corrected to include enhanced classes. If an enhanced class is encountered during path searching, all reads on that path are updated to reflect the enhanced chromosomal information. Section 2 then follows, where reads within each chromosomal cluster are assembled using an existing assembler, and then contigs from all chromosomal bins are subsequently merged to yield the final assembly.

First, complete pangenome graph, which includes nearly all variant sequences of species, with a linear reference genome as the backbone, was employed to mitigate the impact of unclassified reads. Specifically, unclassified reads are aligned to the pangenome graph, and those that align to the linear reference regions are directly assigned to their initial categories based on chromosome IDs. For the reads that align to bubble regions in the pangenome graph, it is difficult to directly determine their classification information due to the lack of chromosome labels in these regions. HiFiCCL addresses this issue by leveraging the pangenome graph structure, assuming that bubble regions with high sequence cohesion to a specific chromosome in the linear reference genome also belong to that chromosome. Specifically, a DFS-based graph traversal algorithm is used, starting from the alignment position of the read in the graph, and performing a bidirectional search until a position in the linear reference genome is reached.

Second, de novo class correction is applied to reads misclassified by the reference genome, and it further reduces the impact of unclassified reads to a greater extent. HiFi reads from the genome to be assembled are used to construct a labeled bidirectional overlap graph, referred to as the Alignment Label Graph (ALG), where each node represents a read, edges represent overlaps, and edge weights indicate the degree of overlap (as defined in the “Materials and Methods”, reflecting the reliability of the connection). Each node is labeled with the chromosome ID to which the read belongs. By starting bidirectional searches from randomly selected nodes, the search is truncated when a significant drop in edge weight is encountered (see “Materials and Methods” section). This process partitions the overlap graph into reliable linear subgraphs, where all nodes within each subgraph are determined to belong to the same chromosome. To mitigate the randomness introduced by the selection of starting nodes, this process is repeated M times with randomized node orders. For each iteration, the chromosome of all nodes (reads) within each subgraph is determined through a voting mechanism. After M iterations, each node’s final class is assigned based on the majority vote across all iterations.

Due to translocation variations between the reference genome and the genome to be assembled, in the reference-based clustering, reads from translocated genomic regions may be assigned to wrong chromosomes. When the translocated region is large, using the above-mentioned correction method may result in a situation where the number of misclassified reads within a subgraph exceeds the number of correctly classified reads, leading to mis-correction. To address this issue, HiFiCCL employs the translocation variation detection algorithm to call translocation variations and the breakpoints in the genome. Based on this, it assigns the reads used for breakpoint identification to their correct chromosome IDs. Then, in the overlap graph, during the search process, these reads are encountered, and all reads within the corresponding subgraph are assigned to the same chromosome as the breakpoint reads.

### Improving contig-level assembly with fewer assembly errors

To evaluate whether the HiFiCCL framework proposed in this study can improve the performance of different existing assemblers on ultra-low coverage HiFi data, we conducted tests on the human HiFi datasets HG002 and NA19240 (∼5x coverage). These tests utilized various assemblers, including Hifiasm (0.19.5-r592), HiFlye (2.9.4-b1799), and LJA (0.2), as the base assemblers for HiFiCCL. These base assemblers were compared and combined with the HiFiCCL using two modes: the primary mode, which only utilizes the CHM13-T2T (CHM13v2.0.fa) (*24*) as a linear reference for clustering guidance, and the optional mode, which employs a human pan-genome reference (*4*) constructed with minigraph (*28*) (hprc-v1.0-minigraph-chm13.gfa). As shown in Table 1, HiFiCCL improved the assembly for Hifiasm, HiFlye, and LJA under ultra-low coverage conditions. Specifically, HiFiCCL-Hifiasm surpassed Hifiasm in nearly all metrics for both HG002 and NA19240 datasets at about 5X coverage, particularly in reducing the misassembled contigs length (MCL). Similarly, HiFiCCL-HiFlye and HiFiCCL-LJA showed better assembly performance compared to their respective base assemblers at about 8X coverage for both datasets. Among these, HiFiCCL-LJA exhibited a relatively significant improvement, especially in further reducing the MCL value, which was already low in the LJA assembly results, while correction strategy in HiFiCCL was highly effective, and it helped mitigate the issues of alignment bias towards linear reference genomes, thereby effectively reducing the size of read sequences that fail to be assigned a chromosome label (Fig. S2).

**Table 1.** Statistics of human primary assemblies. HiFiCCL-assembler indicates the use of different existing assemblers as the base assemblers for HiFiCCL. HiFiCCL represents the primary mode of HiFiCCL and HiFiCCL (optional) represents the optional mode of HiFiCCL. MCL, which stands for “misassembled contigs length”, is a metric used by the quast(29) tool to evaluate the assembly accuracy by comparing assembled genomes against the corresponding high-quality genome. The NG50 of an assembly is defined as the sequence length of the shortest contig at 50% of the total genome size. The NGA50 of an assembly is the length of the correctly aligned block at 50% of the total reference genome size, which is assumed to be 3.1Gb.” Complete” refers to the percentage of all complete single-copy and duplicated orthologous genes. “Single” refers to the percentage of all complete orthologous genes that exist in a single copy. Bolded data indicates that HiFiCCL metric’s performance surpassed that of the base assembler, while a star in the top right corner denotes the best performance achieved.

| Dataset | Assembler | Size (Gb) | Contigs number | MCL (Mb) | NG50 (Kb) | NGA50 (Kb) | Gene completeness (BUSCO)<br>Complete /Single(%) | Missing (%) |
| --- | --- | --- | --- | --- | --- | --- | --- | --- |
| HG002<br>(HiFi 5x) | Hifiasm | 2.73 | 17868 | 331.84 | 194.55 | 174.55 | 82.0/78.7 | 7.6 |
|  | HiFiCCL-Hifiasm | **2.79\*** | 18304 | **274.16** | **199.05\*** | **185.04\*** | **82.6\*/79.1\*** | **7.1\*** |
|  | HiFiCCL-Hifiasm (optional) | **2.79\*** | 18264 | **274.07\*** | **198.76** | **184.97** | **82.6\*/79.0** | **7.1\*** |
| HG002<br>(HiFi 8x) | HiFlye | 3.89 | 46896 | 855.19 | 462.76 | 361.91 | 92.4/81.0* | 2.4 |
|  | HiFiCCL-HiFlye | 3.89 | 46988 | **832.09** | **492.60** | **388.44\*** | **92.9\*/80.9** | **2.1\*** |
|  | HiFiCCL-HiFlye (optional) | **3.88\*** | **46726\*** | **831.81\*** | **493.24\*** | **387.55** | **92.8/81.0\*** | **2.3** |
|  | LJA | 3.09 | 35718 | 120.08 | 148.37 | 138.94 | 78.7/69.4 | 11.1 |
|  | HiFiCCL-LJA | **3.11** | 36083 | **91.84** | **153.38** | **145.13** | **80.5/70.8** | **9.5** |
|  | HiFiCCL-LJA (optional) | **3.11** | 36090 | **90.60\*** | **153.66\*** | **145.54\*** | **80.7\*/71.0\*** | **9.3\*** |
| NA19240<br>(HiFi 5x) | Hifiasm | 2.57 | 22561 | 245.96 | 123.48 | 112.86 | 77.5/74.5 | 10.1 |
|  | HiFiCCL-Hifiasm | **2.64\*** | 23417 | **179.15** | **124.83** | **116.19\*** | **77.9\*/74.9** | **9.9** |
|  | HiFiCCL-Hifiasm (optional) | **2.64\*** | 23405 | **177.83\*** | **125.00\*** | **116.15** | **77.9\*/75.0\*** | **9.8\*** |
| NA19240<br>(HiFi 8x) | HiFlye | 4.11 | 57340 | 777.70 | 333.87* | 267.43* | 91.5/77.5 | 3.0 |
|  | HiFiCCL-HiFlye | **4.10\*** | 57994 | **719.69** | 330.61 | 267.23 | **91.9\*/77.7\*** | **2.8\*** |
|  | HiFiCCL-HiFlye (optional) | **4.10\*** | 57985 | **718.01\*** | 328.83 | 266.08 | **91.9\*/77.5** | **2.8\*** |
|  | LJA | 2.89 | 28917 | 135.50 | 163.10 | 151.39 | 79.2/71.4 | 10.9* |
|  | HiFiCCL-LJA | 2.87 | **28406** | **117.75** | **165.63\*** | **154.14\*** | **79.2/71.5\*** | 11.0 |
|  | HiFiCCL-LJA (optional) | 2.87 | **28365\*** | **115.83\*** | **165.36** | **153.87** | 79.2/71.4 | 10.9* |

HiFiCCL-HiFiasm demonstrated outstanding performance, prompting us to further compare it against other state-of-the-art assemblers, including HiFlye, LJA, Verkko and GALA on the HG002 and NA19240 HiFi dataset at about 5X coverage (Table S1). The results demonstrated that the assemblies produced by HiFiCCL-Hifiasm, Hifiasm, and HiFlye closely approximated the true genome size, whereas those generated by LJA and Verkko were markedly smaller relative to the actual genome size. Among the three methods that produced relatively complete genomes, HiFiCCL-Hifiasm demonstrated similar levels of contiguity and BUSCO completeness but exhibited the lowest misassembly contigs length (MCL). Additionally, it achieved a significantly faster runtime compared to Hifiasm (Table S2), showing that HiFiCCL-Hifiasm not only improves assembly quality but also substantially accelerates the process by more than 1.5 times on ultra-low coverage (∼5X) human datasets, under comparable computational resources and memory usage relative to Hifiasm. It is important to note that we tested GALA with it supported all assemblers, using a linear reference genome and three assembly results as input. However, at ultra-low coverage, due to the generally lower contiguity and higher assembly error rates of input results from other assemblers (such as Hifiasm, HiFlye, LJA) required as input, GALA tended to over-partition the read sequences into significantly more bins than the actual number of chromosomes, ultimately leading to assembly failures. Additionally, GALA’s complexity, particularly the chromosomal clustering step which took over 20 times longer than HiFiCCL (Table S3), may contribute to its unsuitability for ultra-low coverage assembly scenarios.

We further explored the performance of HiFiCCL-Hifiasm on ultra-low coverage HiFi datasets for plants, selecting two species: rice (∼5X) and Arabidopsis thaliana (∼5X). In accordance with the findings from the human datasets, our method demonstrated better performance compared to Hifiasm, achieving the highest BUSCO completeness and the assembly size closely aligned with the true genome size (Table S4). While maintaining comparable contiguity metrics, it also exhibited a significantly smaller misassembled contigs length. Notably, GALA failed to assemble, and its chromosomal clustering time was substantially longer than that of HiFiCCL (Table S5 and S6).

To comprehensively evaluate whether the proposed HiFiCCL demonstrates superior performance over base assemblers in population-scale genome assembly, we compared HiFiCCL-Hifiasm with Hifiasm using 45 human ultra-low coverage HiFi samples (∼5X), consistent with those used in constructing the HPRC draft pan-genome. HiFiCCL utilized the CHM13-T2T as the reference genome for clustering guidance on these datasets. The Quast tool was used to evaluate assembly accuracy by comparing the assembled genomes against the corresponding high-quality reference genomes generated from high coverage sequencing data incorporating multiple data types, as published by the HPRC. As shown in Fig. 2 (Table S7), HiFiCCL-Hifiasm was statistically significantly better than Hifiasm in genome size (Fig. 2A, p<0.0001) and exhibited a statistically significantly lower misassembly contigs length (MCL) (Fig. 2B, p<0.0001), while maintaining comparable contiguity (Fig. 2C, D). Notably, HiFiCCL-Hifiasm reduced the length of misassembled contigs by an average of 21.19% and up to 38.58% relative to Hifiasm. HiFiCCL-Hifiasm also demonstrated statistically significantly higher BUSCO gene completeness (Fig. 2E, p<0.0001), higher single-copy gene completeness (Fig. 2F, p<0.0001), and a lower gene missing rate (Fig. 2G, p<0.0001). These results show that HiFiCCL produces more complete and accurate assemblies with fewer errors and better gene retention at the population-level genome assembly, while also exhibiting excellent generalization capabilities.

**Fig. 2.**
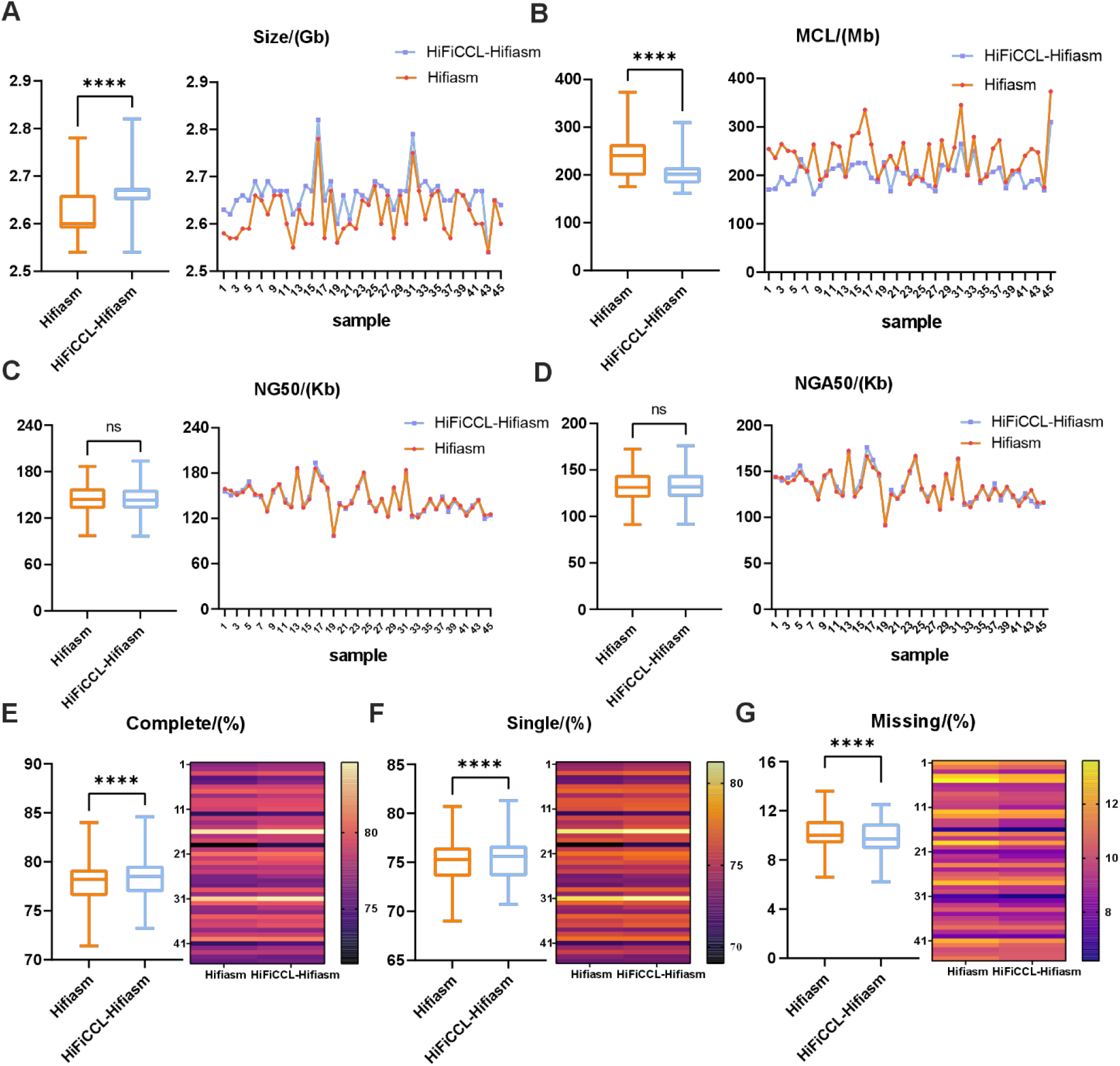
Benchmarking assemblies of Hifiasm and HiFiCCL-Hifiasm on 45 human samples (∼5x coverage). (A) Comparison of assembled genome size metrics between Hifiasm and HiFiCCL-Hifiasm. (B) Comparison of misassemblies contigs length (MCL) metrics. (C) Comparison of NG50 metrics. (D) Comparison of NGA50 metrics. (E) Evaluation of gene completeness metrics. (F) Contrast of single-copy gene completeness metrics. (G) Assessment of missing genome metrics. Statistical significance is indicated as ‘****’ for p < 0.0001, and ‘ns’ denotes no significant difference.

The effect of varying data coverages on the proposed HiFiCCL was assessed by testing it in two modes across different coverage levels of the HG002 dataset (∼3x, 5x, 8x, and 11x), combined with various base assemblers to compare their assembly performance. The results indicated that assemblers based on overlap graph or repeat graph, such as Hifiasm and HiFlye, performed well as base assemblers for HiFiCCL at ultra-low coverages of 3X and 5X (Table S8). As coverage increased, their contiguity metrics significantly improved. And assemblers based on multiplexed de Bruijn graphs like LJA, when used as base assemblers for HiFiCCL, failed to assemble at 3X coverage. At 5X coverage, while the assembled genome size was small and comparable to that of LJA, other metrics were similar, most notably with a significant reduction in the misassembled contigs length. With increased coverage, HiFiCCL-LJA showed notable improvement at 8X coverage, not only surpassing LJA’s assembly performance but also achieving the lowest MCL compared to other assemblers.

### Enhancing assembly-based SVs detection with HiFiCCL’s assemblies

To evaluate the potential of assemblies from HiFiCCL in improving structural variations (SVs) detection, we conducted SVs calling on the assemblies generated by HiFiCCL-Hifiasm and Hifiasm at ∼5X coverage, as well as HiFiCCL-HiFlye, HiFlye, HiFiCCL-LJA, and LJA at ∼8X coverage. The SVs calling was benchmarked against structural variations within the high-confidence regions as defined by the HG002 GIAB Tier 1 dataset (*30*). Firstly, these assemblies were first aligned to the GRCH37 reference genome using minimap2 (*31*), followed by sorting and indexing with samtools (*32*). Secondly, SVs were detected using the assembly-based SV detection tool svim-asm (*33*). The results (Fig. 3, A-C, Fig. S3, Table S9) further validated previous findings, demonstrated that HiFiCCL improved the performance of base assemblers, as evidenced by improved structural variant detection capabilities.

**Fig. 3.**
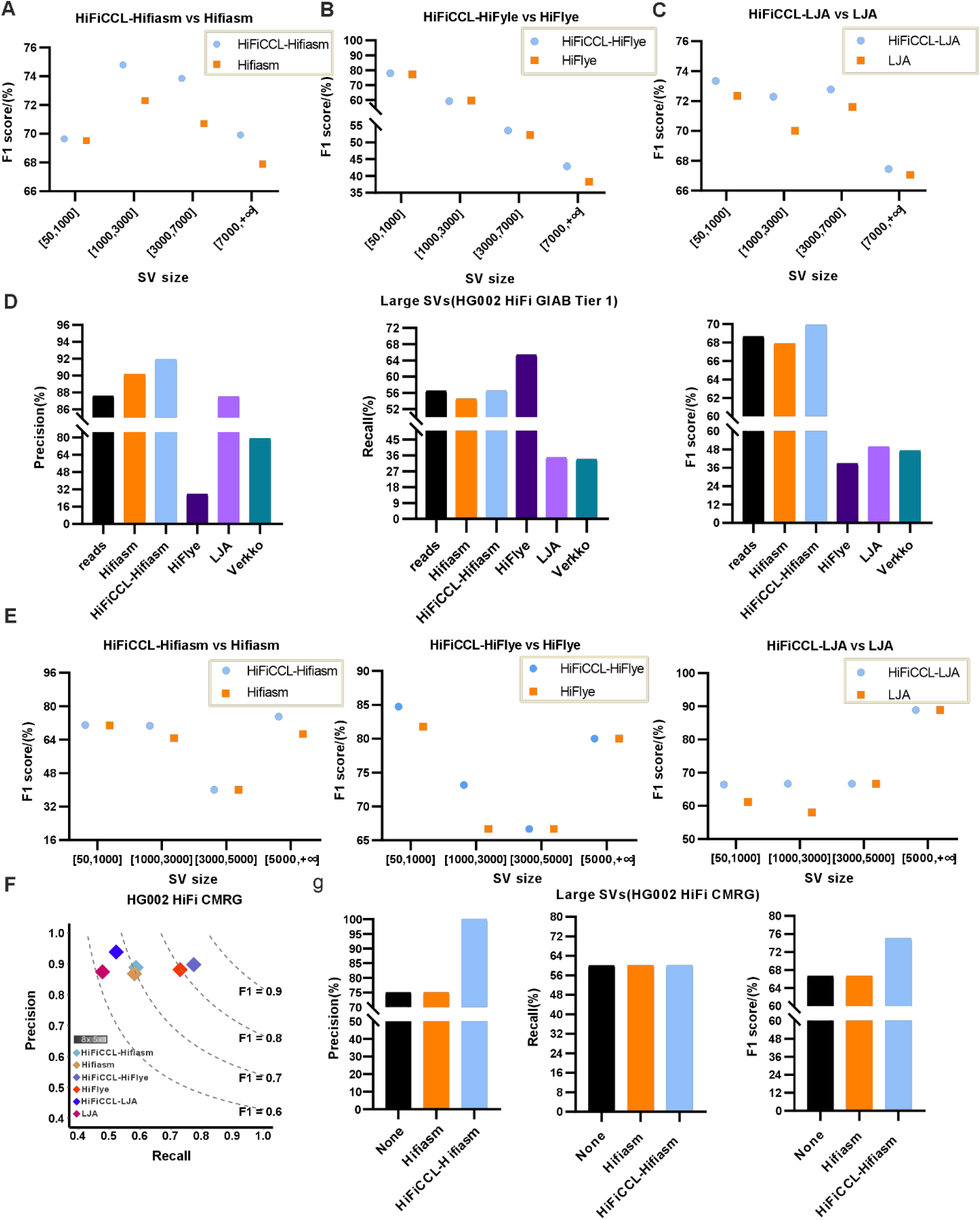
Assembly-based SVs detection and comparison. (A) Comparison of F1 scores for SVs detection (GIAB Tier 1) based on assemblies from HiFiCCL-Hifiasm and Hifiasm across different SVs sizes. (B) Comparison of F1 scores for SVs detection based on assemblies from HiFiCCL-HiFlye and HiFlye at varying SVs sizes. (C) Comparison of F1 scores for SVs detection based on assemblies from HiFiCCL-LJA and LJA across different SVs sizes. (D) Comparison of long SVs detection from various assemblies with the reads-alignment-based SVs calling, where “reads” assemblers represent SVs identified from their respective assemblies. (E) Comparison of SV detection in the CMRG region based on assemblies from HiFiCCL-Hifiasm, Hifiasm, HiFiCCL-HiFlye, HiFlye, HiFiCCL-LJA, and LJA across different SVs sizes. (F) Comparison of SVs detection capability in the CMRG region with precision-recall curves. (G) Comparison of effectiveness in detecting long SVs in the CMRG region using reads-alignment-based and assembly-based SV calling approaches.

Assembly-based SVs detection generally facilitates the identification of larger SVs compared to alignment-based SVs detection (*34*). On the HG002 dataset at 5X coverage, we compared the SVs detection capabilities of the read-alignment-based method svim (*35*) with those of svim-asm, which used alignments from different assemblies (such as Hifiasm, HiFiCCL-Hifiasm, HiFlye, LJA, Verkko). The size distribution of SVs was shown in Fig. S4, where it was evident that assembly-based SVs detection identified more SVs larger than 7000 bp compared to read-based SVs detection. Subsequently, we compared the detection performance for SVs larger than 7000bp (Fig. 3D, Table S10). The results indicated that HiFiCCL-Hifiasm exhibited the highest accuracy and a recall rate comparable to that of the “reads” category (where “reads” is used to denote the reads-alignment-based method for detecting SVs), along with the highest F1 score.

We further evaluated the ability of HiFiCCL’s assemblies in detecting challenging, medically relevant SVs (HG002 CMRG) (*36*), yielding promising results. The results (Fig. 3E, F, Table S11) demonstrated that our method significantly improved the detection of medically relevant SVs compared to the results from base assemblers. The improved detection of medically relevant SVs suggests that HiFiCCL produces more accurate and complete genome assemblies, preserving critical genomic regions that are often challenging to resolve. In addition, based on the commendable performance of HiFiCCL-Hifiasm, Hifiasm assemblies, and reads-alignment-based detection (“reads”) in detecting SVs using the HG002 dataset (5x) within the GIAB Tier 1 benchmarks, we selected these three methods for further evaluation of their efficacy in detecting large, medically relevant SVs (greater than 5000bp) (Fig. 3G, Table S12). The results indicated that HiFiCCL-Hifiasm also performed the best, achieving an accuracy of 1, despite the limited number of SVs in the CMRG set.

### Reducing chromosome-level mis-scaffolding with HiFiCCL’s assemblies

Further scaffolding of the assemblies from HiFiCCL and base assemblers was conducted on ultra-low coverage datasets (HG002 and NA19240) using RagTag (*37*) with CHM13-T2T as the reference genome (Table S13). At the scaffold level, our method displayed similar continuity metrics to the base assemblers but outperformed them in BUSCO completeness and reduced the number of misassemblies (MA). After filtering scaffolds shorter than 5M, we obtained chromosome-level sequences and evaluated baseline metrics, continuity, accuracy, and gene completeness. Consistent with earlier results, HiFiCCL maintained comparable continuity, higher gene completeness, and fewer the number of misassemblies (Table S14).

For synteny analysis, we compared the scaffolding results generated by different base assemblers, both with and without HiFiCCL, against the corresponding reference genomes for each dataset. Specifically, we used the HG002 T2T reference genome (*21*) and the HPRC-released NA19240 assemblies, which were further scaffolded to chromosome level using Ragtag. Synteny analysis was then performed using the NGenomeSyn tool (*38*). The results (Fig. S5, S6) showed a reduction in inter-chromosomal mis-scaffolding with HiFiCCL, particularly at 5X coverage, where HiFiCCL-Hifiasm markedly outperformed Hifiasm. Additionally, using HiFiCCL-Hifiasm as an example, we aimed to investigate its ranking among different assemblers following scaffolding. We scaffolded the assemblies of HiFiCCL-Hifiasm, Hifiasm, HiFlye, LJA, and Verkko from the previously mentioned HG002 (5x) and NA19240 (5x) datasets using Ragtag. The results (Table S15) demonstrated that only the scaffolds from HiFiCCL-Hifiasm, Hifiasm, and HiFlye yielded more complete genomes, echoing previous findings. Among these, HiFiCCL-Hifiasm exhibited the lowest number of misassemblies and the highest completeness of single-copy genes, with other metrics being comparable. This further underscores that our method improves the assembly quality of base assemblers, thereby improving scaffolding performance. After filtering out scaffolding sequences shorter than 5M, our method continued to exhibit the lowest number of misassemblies (MA) at the chromosomal level (Table S16). We then conducted a synteny analysis of these scaffolds (Fig. 4, Fig. S7). It was observed that the chromosome-level scaffolds of HiFiCCL-Hifiasm post-scaffolding exhibited the best performance, with inter-chromosomal mis-scaffolding comparable to those of Verkko post-scaffolding, while achieving a more complete genome size.

**Fig. 4.**
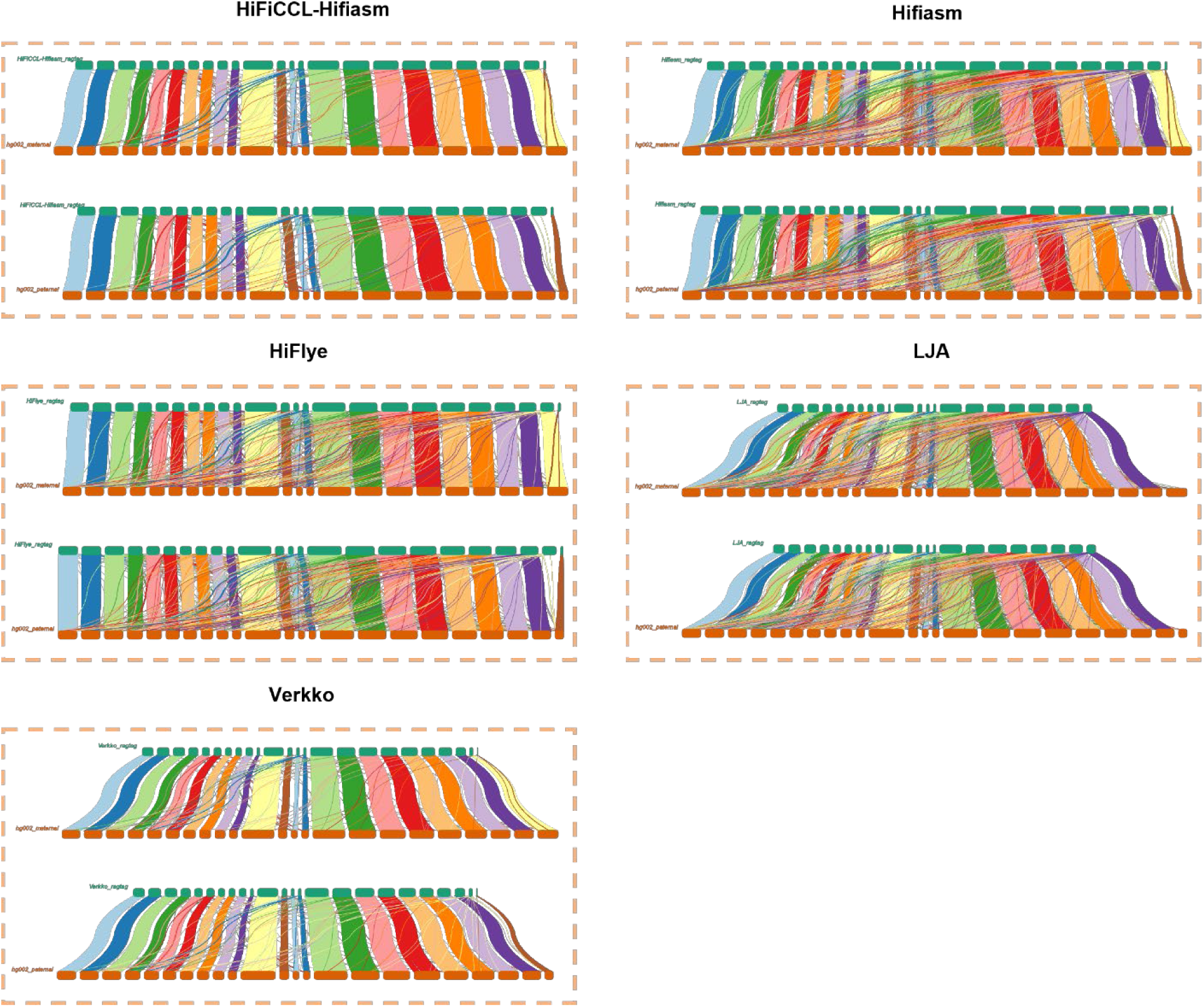
Synteny analysis of scaffolding results on ultra-low coverage HiFi dataset for HG002. It shows the synteny between the scaffolding performance of different assemblies and the maternal and paternal reference genomes for HG002. Within each box, the top panel represents the synteny analysis with the maternal reference genome, while the bottom panel represents the synteny analysis with the paternal reference genome.

We further conducted a synteny analysis across six human datasets against their corresponding chromosome-level assemblies, which were generated from HPRC-released assemblies further scaffolded using Ragtag. Four of these datasets (HG00438, HG00621, HG00673, HG00735) had previously shown significant reductions in MCL with HiFiCCL compared to Hifiasm. The other two datasets (HG005, HG01109), which showed only marginal improvements in previous experiments, were included to investigate whether HiFiCCL could achieve better results after scaffolding to the chromosome level. We used Ragtag to scaffold the assemblies of HiFiCCL-Hifiasm (5x) and Hifiasm on these human HiFi datasets and subsequently performed synteny analysis. The results further demonstrate that HiFiCCL-Hifiasm-based scaffolding significantly reduced inter-chromosomal mis-scaffolding across all six datasets compared to Hifiasm-based (Fig. 5, Fig. S8).

**Fig. 5.**
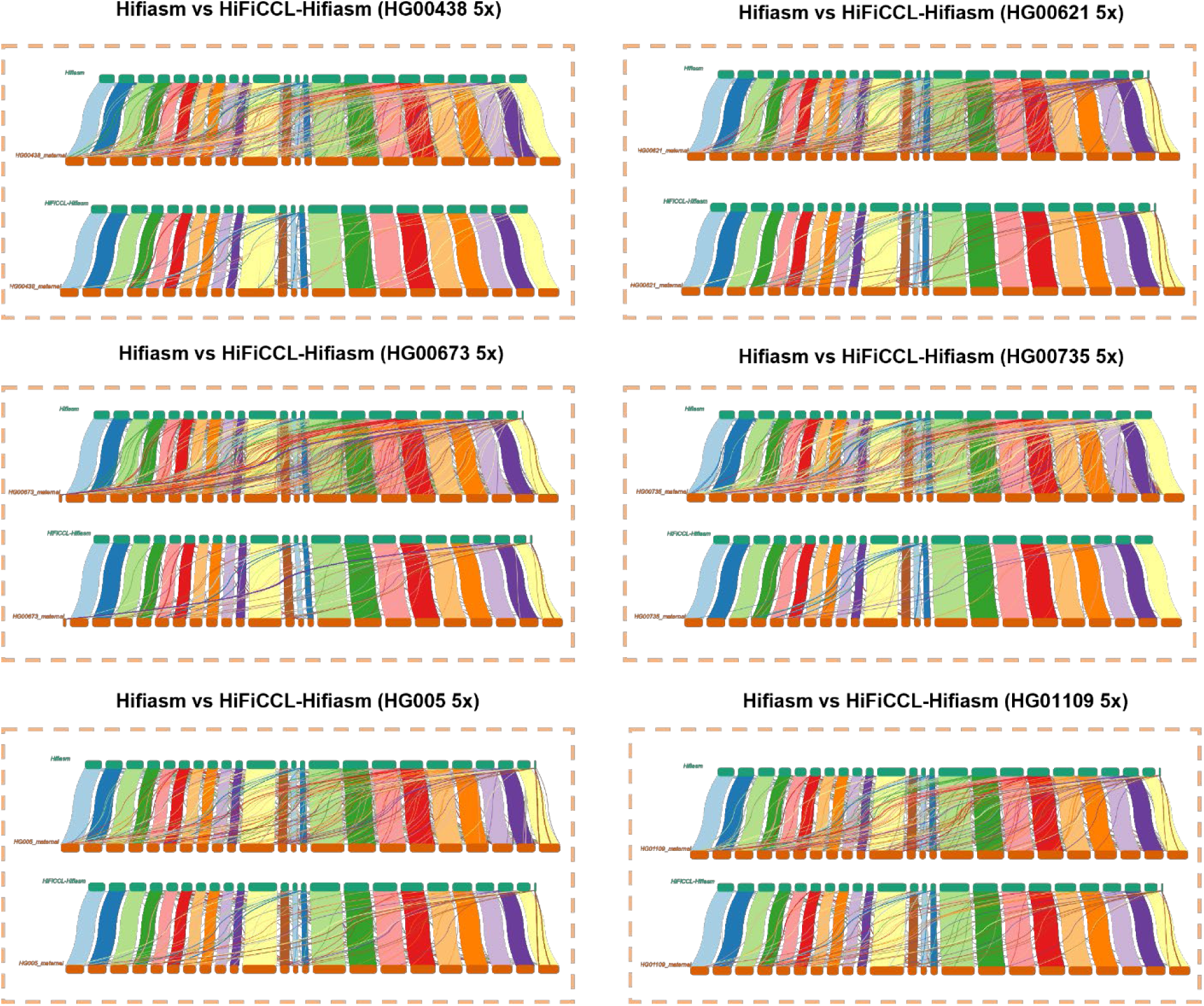
Synteny analysis of scaffolding results across multiple human datasets. In each box, the top section shows the synteny analysis based on the base assembler, while the bottom section displays the synteny analysis after incorporating HiFiCCL. It shows the scaffolding performance of HiFiCCL-Hifiasm and Hifiasm assemblies, with the maternal reference genome used as the reference, across the HG00438, HG00621, HG00673, HG00735, HG005 and HG01109 datasets (∼5x).

### Accurately constructing pangenome graph

To explore the potential applications of HiFiCCL at the population genome level, we constructed human pangenome graphs and evaluated their consistency with the HPRC-released real pangenome graph. Using minigraph, we built Hifiasm_graph with GRCH38, CHM13-1.0, and Hifiasm assemblies from 45 human HiFi datasets (∼5X). Similarly, HiFiCCL-Hifiasm_graph was constructed using the same reference genomes and inputs but with HiFiCCL-Hifiasm assemblies from the same datasets. To evaluate the accuracy of the pangenome graphs, we compared the consistency of bubble regions between Hifiasm_graph and HiFiCCL-Hifiasm_graph against real_graph, built by the HPRC using the same datasets. Bubble region consistency is a key metric that reflects how closely a pangenome graph represents the true population pangenome structure, with higher consistency indicating greater accuracy (see “Materials and Methods” section). As shown in Fig. 6A, HiFiCCL-Hifiasm_graph demonstrated greater consistency and improved accuracy in the bubble regions compared to Hifiasm_graph, suggesting that HiFiCCL assemblies generate pangenome graphs that more closely approximate the real pangenome. This result highlights the utility of HiFiCCL in improving pangenome construction and its potential for advancing population-level genomic studies.

**Fig. 6.**
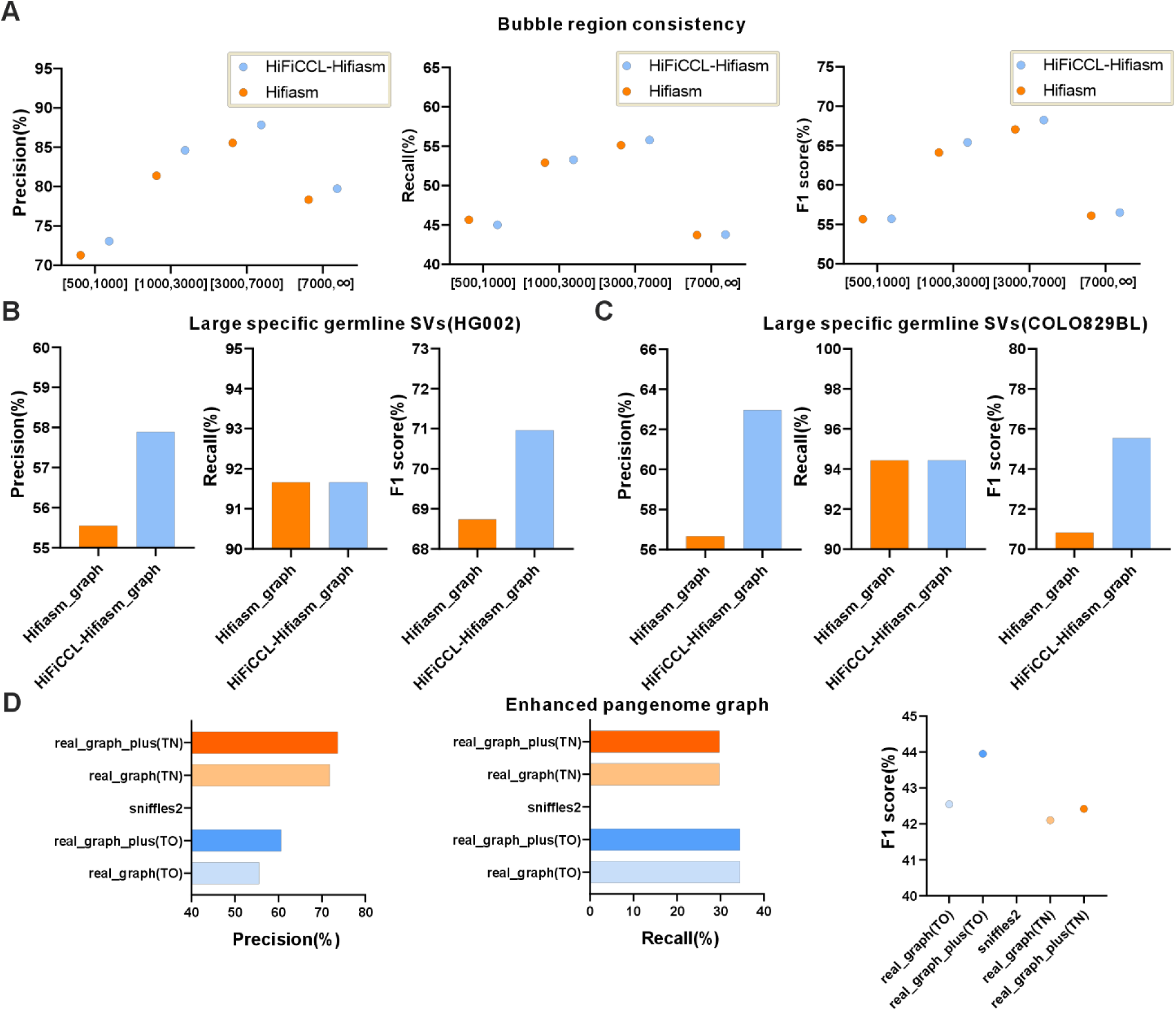
Pan-genome graph consistency analysis and applications. (A) The consistency of bubble regions between pan-genome graphs constructed using HiFiCCL-Hifiasm and Hifiasm assemblies from 45 human samples with GRCh38 and CHM13 sequences against the pan-genome graph released by the HPRC. (B) Detection of large-rare germline SVs in the HG002 dataset using pan-genome graphs constructed from Hifiasm or HiFiCCL-Hifiasm assemblies. (C) Detection of large-rare germline SVs in the COLO829BL dataset using pan-genome graphs constructed from Hifiasm or HiFiCCL-Hifiasm assemblies. (D) Detection of cancer somatic SVs using the pan-genome graph released by the HPRC (real_graph) and its enhanced version, real_graph_plus, where TO indicates the use of only tumor tissue datasets, TN indicates the use of both tumor and normal tissue datasets, and Sniffles2 was used solely for detection using the tumor dataset.

### Facilitating the detection of Large-rare germline SVs using pangenome graph

The constructed pangenome graphs can serve as a powerful tool for filtering common germline SVs, thereby enabling the detection of large, rare germline SVs. Therefore, we detected large-rare germline SVs (>=10,000bp) in HG002 and COLO829BL datasets based on pangenome graphs constructed from different assemblies. SVs detection was performed using minisv (https://github.com/lh3/minisv), a tool developed by Li Heng’s team, specifically designed for SVs detection on pangenome graph. For the HG002 HiFi dataset (∼37X), alignments were first performed against GRCh38 and the CHM13-T2T reference genomes using minimap2. Subsequently, the dataset was aligned to the Hifiasm_graph, HiFiCCL-Hifiasm_graph, and real_graph separately using minigraph. The alignment results from both linear references and different graphs were input into minisv. SVs detected based on the real_graph alignments were used as the ground truth to assess the performance of Hifiasm_graph-based and HiFiCCL_graph-based. As shown in Fig. 6B, the HiFiCCL_graph-based method had the higher accuracy with comparable recall to the Hifiasm_graph-based approach. Similar results were observed on the COLO829BL nanopore dataset (∼30X) (Fig. 6C).

### Facilitating the detecting tumor somatic SVs using pangenome graph

Somatic SV mutations in tumors often involve large structural variations, making their detection crucial for understanding tumor genomics, guiding personalized treatments, and drug development (*39*). The minisv offers two modes for detecting cancer somatic SVs: tumor-only and tumor-normal paired modes. In the tumor-only mode, the COLO829 nanopore dataset (∼30X) (*39*) was aligned separately to GRCH38, CHM13-T2T, “real_graph”, “Hifiasm_graph”, and “HiFiCCL-Hifiasm_graph”. The results from different graph alignments, combined with those from alignments to the two linear reference genomes, were respectively input into minisv for the detection of cancer somatic SVs larger than 10,000bp. Evaluation against the COLO829 standard SV set (*40*) (>=10,000 bp) showed that HiFiCCL-Hifiasm_graph-based achieved higher accuracy than Hifiasm_graph-based with the same recall (Fig. S9). In the tumor-normal paired mode, both COLO829 and COLO829-BL (∼30X) datasets were utilized. HiFiCCL-Hifiasm_graph-based outperformed Hifiasm_graph-based in accuracy and matched real_graph in accuracy, recall, and F1 score (Fig. S9).

We further assembled additional 20 human samples using HiFiCCL-Hifiasm and enhanced “real_graph” with these assemblies to create “real_graph_plus”. The COLO829 dataset (∼30x) was aligned to “real_graph_plus”, and the alignment results, along with those from two linear reference genomes, were analyzed using minisv in tumor-only mode. As shown in Fig. 6d, SV detection (over 10,000 bp) based on “real_graph_plus” significantly improved accuracy compared to “real_graph”, with no change in recall. This suggests that the assembly results from HiFiCCL at ultra-low coverage can enhance “real_graph”, improving large somatic SV detection in cancer. In the tumor-normal paired mode, incorporating cross-alignment information between tumor and normal datasets allowed for a more comprehensive analysis. We compared cancer somatic SVs larger than 50 bp using real_graph-based and real_graph_plus-based approaches. The results demonstrated that the real_graph_plus-based method significantly improved detection precision. Moreover, Sniffles2 (*41*) failed to detect tumor somatic SVs when only tumor sample dataset (∼30x) was available.

## Discussion

Second-generation sequencing in population genomics is limited by its inability to resolve complex genomic regions, such as structural variations and repetitive sequences. The advent of third-generation sequencing, particularly HiFi data, has propelled the development of telomere-to-telomere (T2T) reference sequences and pangenome, marking a new era in genomic research with unprecedented resolution and representation of genetic diversity. However, while PacBio HiFi reads offers high accuracy and long-read capabilities, its high cost and the poor performance of most assemblers at ultra-low coverage hinder its application in large-scale studies. To address these challenges, we developed HiFiCCL, a reference-guided chromosome-by-chromosome assembly framework that leverages high quality references or pangenome graph for efficient chromosomal clustering and assembly with ultra-low coverage HiFi reads.

With the release of an increasing number of telomere-to-telomere (T2T) and pangenome sequences for various species, HiFiCCL, leveraging its reference-guided assembly capabilities, holds significant potential and advantages for large-scale population genomic studies at ultra-low coverage condition. HiFiCCL offers the ability to process more samples with reduced costs and computational requirements, making it highly suitable for preliminary studies on population genetic structure, evolutionary relationships, and genetic diversity. It can also facilitate the screening and prioritization of samples or genomic regions for more in-depth analyses. Integrating additional data types, such as Hi-C or nanopore sequencing, could further enhance chromosome-by-chromosome assembly performance. Our future efforts will focus on adapting this approach to support T2T sequence assembly at lower coverage, enabling cost-effective high-precision genetic analyses.

In summary, HiFiCCL innovatively leverages high quality reference sequences or pangenome graphs to guide chromosomal clustering of read sequences and implements a chromosome-by-chromosome assembly strategy. This approach has consistently demonstrated strong performance across key areas, including improving assembly quality in ultra-low coverage HiFi datasets, improving the detection of structural variations (SVs), minimizing inter-chromosomal mis-scaffolding, and delivering reliable results in population genomics studies. As more T2T reference sequences and pangenome assemblies become available, and as pangenome tools are further refined, HiFiCCL is expected to unlock even greater potential. Moreover, its applicability extends to large-scale ultra-low coverage human genome assemblies, making it a promising tool for studies investigating host genetics and gut microbial genetic diversity (*42–44*).

## Materials and Methods

### Construction of Alignment Label Graph

HiFiCCL utilizes two modes for obtaining initial chromosome classes for reads: the primary linear reference genome guided mode and the optional pan-genome guided mode. In the primary mode, the initial chromosome classes for reads are determined by aligning sequences with the high-quality linear reference genome using the alignment tool minimap2, with parameters set to “-ax map-hifi”. In the optional mode, reads that fail to align in the primary mode are assigned chromosome classes by aligning them to the pangenome graph using minigraph with parameters set to “-cx lr”. Subsequently, minimap2 is used to perform inter-sequence comparisons for all read sequences with parameters set as “-D --dual=no --no-long-join -k19 -w5 -U50,500 --rmq -A1 -B19 -O39,81 -E3,1 -H -e0 -m100 -N20”. This process identifies the best alignment position for each sequence relative to others, excluding itself, and identifies multiple suboptimal alignment positions.

Based on the inter-sequence comparison information, a bidirectional graph is constructed as shown in Fig. 1. In this graph, nodes represent the IDs of each sequence, and the edges between nodes indicate the degree of overlap between sequences. The overlap degree is calculated according to equation (1).

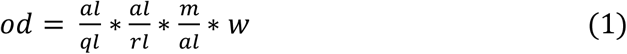

Which *od* denotes the overlap degree between sequences, *al* represents the alignment length between sequences, *ql* signifies the length of the query sequence, *rl* indicates the length of the reference sequence, *m* refers to the length of the match, and *w* is a constant. It is noteworthy that during the graph constructing process, if two sequences exhibit a containment relationship, the node representing the shorter sequence is collapsed into the node of the longer sequence. When analyzing inter-sequence comparison data, if a node *v* align with another node *w*, and *v* and *w* do not share a containment relationship, a directed forward edge *⍺* is established from *v* to *w* with a weight equal to the overlap degree between *v* and *w*. Concurrently, a reverse edge *β* from *w* to *v* is constructed, carrying the same weight, representing the overlap degree between *w* and *v*. Chromosome classes, derived from the alignment of sequences with the reference sequence, are assigned to each node within the graph. For sequences associated with multiple chromosome classes, all such classes are aggregated into the sequence. If a sequence fails to align to the reference genome, its chromosome class is designated as “None”. Utilizing these two types of alignment information, the construction of the final ALG is completed.

### Construction of Pangenome Label Graph

In the optional mode of HiFiCCL, it is necessary to realign reads that failed to align with the linear reference sequence in the primary mode to a pan-genome graph to ascertain the chromosomal classes of these reads. Given that the chromosomal information of the linear reference genome does not coincide with that of the pan-genome graph, a PLG (Pangenome Label Graph) is constructed to harmonize the chromosomal classes. This graph is based on the paths of the pan-genome graph, with nodes representing the nodes of the pan-genome graph, each tagged with its ID. Bidirectional edges are established based on the paths of the classes beginning with “NA” or “HG.” These reads are processed through bidirectional DFS (Depth-First-Search) within the constructed PLG until chromosomal classes starting with “chr” are encountered. After standardizing the chromosomal classes, the chromosomal classes of the nodes in the ALG are assigned accordingly.

### *De novo* class correction

Through the construction of the ALG, we have obtained a graph containing inter-sequence alignment information along with chromosomal classes. The initial chromosomal classes of sequences are “None” if the sequences fail to align with the reference genome. Additionally, similar genome regions across different chromosomes can also cause classing errors. Inspired by the Z-drop algorithm(31), we propose a heuristic algorithm to search the ALG and correct sequence classes, thereby completing the classes correction of reads.

Let *R* = {*r*_1_, *r*_2_, …, *r*_*n*_} denote the set of reads, where *n* is the total number of reads (nodes in the graph). Let *G*(*V*, *E*, *W*) represent the Alignment Label Graph (ALG), where *V* is the set of nodes (reads). *E* is the set of edges representing overlaps between reads. W(*e*_*ij*_) is the weight of the edge between nodes *i* and *j*, reflecting the degree of overlap between reads *r*_*i*_ and *r_*j*_*. Define *C*(*v*_*i*_) as the chromosomal class of node *v*_*i*_ (initially assigned based on alignment to a reference genome or “None” if unclassified). A node list *Nodelist* is created from the set of all nodes *V* in the graph. The order of *Nodelist* is randomized to ensure unbiased traversal in each iteration. Nodes in *Nodelist* are traversed sequentially in the randomized order. For each node *v*_*i*_, a bidirectional search (forward and reverse) is performed to construct a path *PATH*_*k*_, representing a linear subgraph. The path is extended by following a set of rules: only unvisited neighboring nodes are considered for traversal, and among multiple edges, those connected via higher-weight edges are prioritized using a greedy algorithm. If a node to be visited has already been visited, the search continues along the next priority level of edges until there are no accessible nodes left or until a node satisfies equation (2) (indicating a significant drop in overlap degree). Nodes that do not satisfy equation (2) are added to the path *PATH*_*k*_. Equation (2) is described as follows:

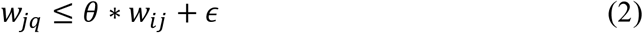

Which *w*_*jq*_ represents the weight of the edge from the current node *j* to the newly accessed node *q*, *θ* is a fraction less than 1, *w*_*ij*_ denotes the weight of the edge from the previous node *i* to the current node *j*, and *∈* is a constant.

For each path *PATH*_*k*_, the chromosomal class is updated based on the majority vote among the nodes in the path:

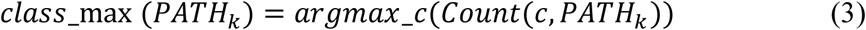

Where *Count*(*c*, *PATH*_*k*_) is the number of occurrences of chromosomal class *c* in *PATH*_*k*_. *class*_max (*PATH*_*k*_) is the most frequently occurring chromosomal class in the path. If the number of nodes in the path, *N* (*PATH*_*k*_), satisfies the minimum support threshold *N*(*PATH*_*k*_) > *SN*_*min*, the *class*_max (*PATH*_*k*_) is propagated as the updated class for all nodes in the path. All nodes in the path are marked as visited (visited =1). The updated chromosomal classes are stored in a new hash table *N*o*deHash*, with the node ID as the key and its current class as the value. Once all node *v*_*i*_ ∈ *V* are traversed, a single iteration is complete. The chromosomal class information from this iteration is stored in a separate hash table *ClassHash*, where:

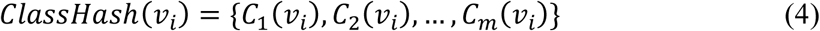

*C*_*k*_(*v*_*i*_) is the chromosomal class assigned to node *v*_*i*_ in the *k*-th iteration. The above process is repeated *M* times, with a new randomized order of *Nodelist* in each iteration. Each iteration generates a set of chromosomal class assignments for all nodes, which are recorded in *ClassHash*. After *M* iterations, the final chromosomal class of each node *v*_*i*_ is determined by majority voting across all iterations:

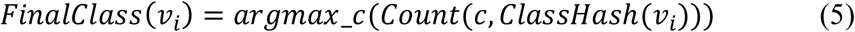

Where *Count*(*c*, *ClassHash*(*v*_*i*_)) is the number of times chromosomal class *c* appears in the list *ClassHash*(*v*_*i*_) over *M* iterations. *FinalClass*(*v*_*i*_) is the most frequently occurring chromosomal class for node *v*_*i*_. Finally, reads are clustered into chromosomal bins based on their final assigned classes:

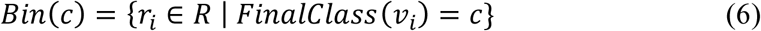

Where *Bin*(*c*) represents the set of reads belonging to chromosomal class *c*.

### Translocation class correction

Translocation variations between the input reads and the reference genome may lead to erroneous classes. Based on the alignment information with a linear reference sequence, translocation variant detection aims to identify the breakpoints where translocation occurs. This enables the correction of reads containing breakpoints information. The translocation detection module primarily employs methods like those used in the translocation detection section of CuteSV(*45*). When a sequence exhibits split alignments during the matching process, each split alignment is recorded as a 6-tuple:

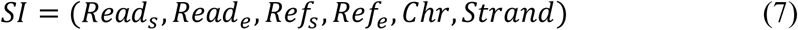

where *SI* represents the split alignment information, *Read*_*s*_ denotes the start coordinate of the aligned region in the read, *Read*_*e*_ denotes the end coordinate of the aligned region in the read, *Ref*_*s*_ signifies the start coordinate of the aligned region in the reference sequence, *Ref*_*e*_ marks the end coordinate of the aligned region in the reference sequence, *Chr* specifies the chromosome information, and *Strand* indicates the direction of the alignment. If two segments of a read are aligned to different chromosomes and the distance between these segments on the sequence is less than 100bp, their split alignment information is extracted as a translocation variant signal, as demonstrated in equation (8).

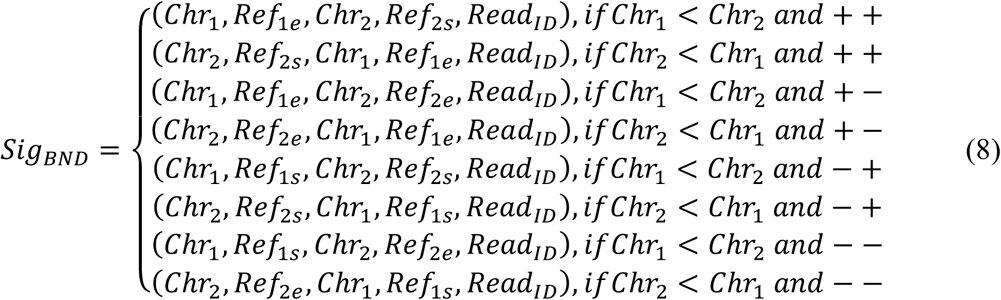

Wherein “<” indicates that the first chromosomal label is alphabetically less than the second chromosomal label, and the combinations “++”, “+-”, “-+”, “--” represent the orientations of structural variations between chromosomes.

Subsequently, the *Sig*_*BND*_ information is clustered by first sorting it based on genomic coordinates and creating a cluster. All signals are scanned from top to bottom to determine if they satisfy equation (9). If a signal meets the criteria, it is added to the cluster. If it does not, a new cluster is created, and the scanning continues to assess whether additional signals can be added to an existing cluster. This process is repeated until all signals are assigned to clusters. Each cluster is then evaluated to determine if the number of signals it contains meets a predefined threshold. Clusters that do not meet this threshold are discarded. The remaining clusters are further clustered using the aforementioned method to identify three breakpoints associated with translocation variants, including two deletion breakpoints and one insertion breakpoint. Equation (9) is described as follows:

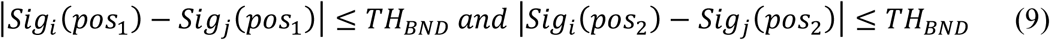

Where *Sig*_*i*_(*pos*_1_) represents the prior coordinate information of signal *i*, and *Sig*_*j*_(*pos*_1_) represents the prior coordinate information of signal *j*. *TH*_*BND*_ is the predefined threshold, *Sig*_*i*_(*pos*_2_) denotes the subsequent coordinate information of signal *i*, and *Sig*_*i*_(*pos*_2_) denotes the subsequent coordinate information of signal *j*. The labels of reads carrying translocation breakpoints signals are assigned to the chromosomal information of the insertion breakpoint site and designated as enhanced nodes. This designation implies that, during *de novo* correction, if these reads are encountered, the classes of all reads in the path will be assigned to the chromosomal information represented by the enhanced nodes.

### A chromosome-by-chromosome assembly strategy

By clustering input reads based on their respective chromosomes’ information, the original problem of assembling mixed chromosomal reads is transformed into assembling reads for each individual chromosome, significantly reducing the complexity of the original assembly process. This approach is particularly advantageous in ultra-low coverage datasets, where insufficient sequence information during the path selection process can lead to misassemblies between chromosomal sequences. Clustering reads by chromosome helps mitigate this issue by reducing the likelihood of inter-chromosomal misassemblies under such challenging conditions. Moreover, this allows for the parallel assembly of each chromosomal data, accelerating the assembly process. The specific workflow is as follows: First, utilize inter-sequence alignment information and sequence-to-reference alignment information to construct an ALG. Next, use *de novo* class correction and translocation class correction to correct the chromosomal classes of the reads, and cluster the sequences according to their respective chromosomes. Assemble the reads for each chromosome using an existing assembler. Finally, merge the assemblies from each chromosome bin to form the final assembly.

### Processing of reads and evaluation of assemblies

These reads were down sampled using the seqtk tool (1.3-r106). To evaluate the assemblies, we utilized QUAST (v5.2.0) to obtain basic information about the assembled contigs and assess their continuity and assembly accuracy, including contigs size, number of contigs, misassembled contigs length, as well as NG50 and NGA50 metrics. Accuracy metrics, such as misassembled contigs length (MCL) or the number of misassemblies (MA), were evaluated using high-quality published assemblies corresponding to ultra-low-coverage datasets. BUSCO (5.4.7) (*46*) was employed to assess gene completeness, using the database of conserved single copy orthologs for vertebrates in human genomes. For rice and Arabidopsis thaliana genome evaluation, the database of conserved single copy orthologs for angiosperms was utilized. All experimental commands related to this study are provided in Supplementary Text.

### Detection and evaluation of germline SVs

SV detection based on reads alignment was performed using SVIM (2.0.0), while assembly-based SV detection was conducted using SVIM-asm (1.0.3). The evaluation was carried out with Truvari (v3.2.0) (*47*), using GRCh37 as the reference genome and the HG002 GIAB Tier 1 high-confidence regions as the SVs benchmark set. The Challenging Medical Relevant Genes (CMRG) SV panel was also utilized for the assessment.

### Scaffolding, evaluation, and synteny analysis of assemblies

The scaffolding of different assemblies was performed using the Ragtag tool, with CHM13 (v2.0) selected as the reference genome. The evaluation of the scaffolding results was conducted using Quast and BUSCO. When using Quast for evaluation, CHM13 (v2.0) was selected as the reference genome for assessing contiguity metrics, and the reference genome for accuracy metrics was the sequence scaffolded from the high-quality assemblies corresponding to the ultra-low coverage datasets. Synteny analysis was performed using NGenomeSyn.

### Pangenome graph construction and bubble region consistency evaluation

First, the down sampled HiFi (5x) datasets from 45 human samples were assembled using both HiFiCCL-Hifiasm and Hifiasm, respectively, with each sequence renamed to ensure uniqueness across all assemblies. Then, using the GRCh38 sequence as the reference, the input data included the CHM13 (v1.0) sequence along with the HiFiCCL-Hifiasm or Hifiasm assemblies for each sample, sorted alphabetically. These were used to construct the pangenome graph. The bubble regions of the pangenome were identified using gfatools, which generated a BED file representing these regions. Using the BED file generated from the pangenome graph constructed by minigraph from the HPRC release as the reference, bubble regions in our pangenome graph with over 80% overlap with the reference were considered true positives (TP). Bubble regions that did not meet this overlap threshold were classified as false positives (FP), while bubble regions in the reference that did not overlap by at least 80% with our graph were considered false negatives (FN). The evaluation was conducted across different intervals according to the following formulas:

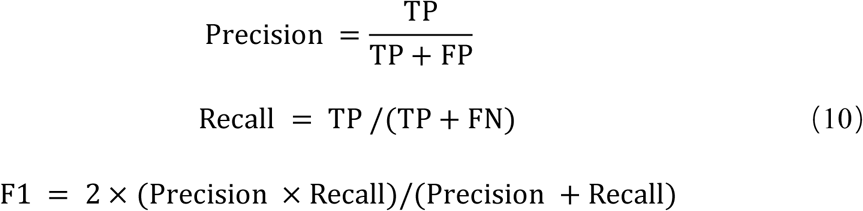

### Large, rare germline SVs detection based on pangenome graph

The input dataset was first aligned to the reference genome GRCh38 using min-imap2, followed by alignment to the CHM13-T2T reference genome using the same tool. Subsequently, the input dataset was aligned to Hifiasm_graph, HiFiCCL-Hifiasm_graph, and real_graph using minigraph. The alignment results from the two linear reference genomes, as well as the alignment to Hifiasm_graph, were then input into minisv for SV detection. Similarly, SV detection results were generated by minisv based on the alignments to the different graphs. The SV detection results from the alignment to “real_graph” were considered the ground truth and used to assess the performance of SV detection for variants larger than 10,000 bp, using the alignment results from “Hifiasm_graph” and “HiFiCCL_graph” as inputs.

### Cancer somatic SVs detection based on pangenome graphs

The minisv supports two scenarios for cancer somatic SVs detection: tumor-only and tumor-normal paired. In the tumor-only scenario, the tumor dataset COLO829 (30X) was aligned to the reference genomes GRCh38, CHM13-T2T, and real_graph (following the same procedure as in large, rare germline SV detection), and the results were input into minisv to obtain the SVs results using the real_graph alignment. Similarly, SVs results were respectively obtained using the alignments to Hifiasm_graph and HiFiCCL-Hifiasm_graph as inputs. The SVs larger than 10,000 bp were then evaluated using the COLO829 SV benchmark set.

In the tumor-normal paired scenario, the input for the three aforementioned minisv alignments was supplemented with reciprocal alignment information between the COLO829 dataset and the COLO829BL dataset (30X) using minimap2.

### Pangenome graph augmentation

Twenty human samples were assembled using HiFiCCL-Hifiasm, and minigraph was then used to augment the “real_graph”, incorporating additional variation and im-proving the representation of pangenomic structures.

### Statistical Analysis

We used the non-parametric Wilcoxon test for statistical analysis, where “****” indicates p < 0.0001, and “ns” denotes no significant difference.

## Acknowledgments

We sincerely thank Chonghui Liu, Hongfei Li, Qianzi Lu, and Xin Lu for their valuable suggestions and insightful discussions that contributed to this study, as well as the opinions and suggestions from other members of the research group.

## Funding

This work has been supported by:

National Key R&D Program of China grant 2022YFF1202101

National Natural Science Foundation of China grant 62225109

## Author contributions

Conceptualization: ZJ, WP, GW

Methodology: ZJ, WP, GW

Investigation: ZJ, WP, GW, RG, HH, WG, ZQ, SJ

Visualization: MZ, YY

Supervision: GW, WP

Writing—original draft: ZJ, WP, GW

Writing—review & editing: ZJ, WP, GW, MZ, YY

## Competing interests

The authors declare that there is no conflict of interest regarding the publication of this paper.

## Data and materials availability

All data are available in the main text or the supplementary materials. The human HiFi data were obtained from the NCBI Sequence Read Archive: two runs (SRR10382244, SRR10382245) for HG002, SRR14611231 for NA19240. The CHM13v2.0 reference generated by the T2T consortium can be found at https://github.com/marbl/CHM13. The HG002 T2T reference can be found at https://github.com/marbl/hg002. The human pangenome graph constructed with CHM13+Y used as reference sequences can be found at https://github.com/human-pangenomics/hpp_pangenome_resources?tab=readme-ov-file. The 45 human samples, along with 20 additional samples utilized for graph augmentation, were all sourced from the Human Pangenome Project accessible at https://s3-us-west-2.amazonaws.com/human-pangenomics/index.html?prefix=working/. The rice HiFi data (GSA: CRR573321) were obtained from the National Genomics Data Center database (NGDC) under project accession number PRJCA012143. The assembly for CX20 rice can be found under project accession PRJCA012309 in NGDC. The genome assembly for T2T-NIP is deposited in the National Center for Biotechnology Information database under project accession number PRJNA953663 and the National Genomics Data Center database under project accession number PRJCA018610. The Arabidopsis thaliana HiFi data (GSA: CRA004538) were obtained from the NGDC. The near complete genome of Arabidopsis thaliana is available under the NGDC accession PRJCA007112. The high-quality Arabidopsis thaliana genome assembly for assembly evaluation can be found in the Genome Warehouse at the NGDC (GWH: GWHBDNP00000000.1). For BUSCO, the embryophyta and vertebrata datasets are available at https://busco-data.ezlab.org/v5/data/lineages/. HiFiCCL code is available at https://github.com/zjjbuqi/HiFiCCL.

## Supplementary Materials

Supplementary Text

Figs. S1 to S9

Tables S1 to S16

## Notes

### Competing Interest Statement

The authors have declared no competing interest.

