## Supplemental information for "Reference-guided genome assembly at scale using ultra-low-coverage high-fidelity long-reads with HiFiCCL"

Zhongjun jiang *et al.*

**This PDF file includes:**

Supplementary Text
Figs. S1 to S9
Tables S1 to S16

**Supplementary Text**

Commands used by different assemblers and evaluation for human and plants datasets

The reference genome used for HiFiCCL is CHM13-T2T (v2.0), while the pangenome utilized is hprc-v1.0-minigraph-chm13.gfa.

HiFiCCL: > python hificcl.py -t 20 -o <your_path> -m n -r <your_path/chm13v2.0.fa> -f <your_input.fasta> -a <assemblers>
HiFiCCL(optional)：> python hificcl.py -t 20 -o <your_path> -m p -r <your_path/chm13v2.0.fa> -R <your_path/hprc-v1.0-minigraph-chm13.gfa> -f <your_input.fasta> -a <assemblers>
Hifiasm：> hifiasm -o <your_path> -t 20 --primary <your_input.fasta>
LJA：> lja -t 20 --diploid -o <your_path> --reads <your_input.fasta>
Verkko：> verkko -d <your_path> --hifi <your_input.fasta> --threads 20
Flye: > flye --pacbio-hifi <your_input.fasta> --out-dir <your_path> --threads 20 --iterations 0
GALA: > gala <your_draft_genome> fa <your_input.fasta> pacbio-corrected --hifi -a <assemblers> -threads 20

In the case of GALA, it is necessary to specify the path to the draft genome. The draft genomes selected are the assembly results from Hifiasm, HiFlye and CHM13(2.0). The code from the original GALA paper available at https://github.com/ganlab/GALA could not be executed successfully. Instead, we used the code from https://github.com/JohnUrban/GALA for our analysis.

The commands used for the evaluation of the assembly results are as follows:
BUSCO: > busco -i <your_input.fasta> -o <your_prefix> -c 20 vertebrata_odb10 --mode genome -f --offline

For plant genomes:
BUSCO: > busco -i <your_input.fasta> -o <your_prefix> -c 20 embryophyta_odb10 --mode genome -f --offline
Quast: > python quast.py -o <your_path> -r <your_reference.fasta> -t 20 <your_input.fasta>

For evaluating basic metrics like contiguity in human datasets, the reference genome used was CHM13 (v2.0). The assembly accuracy was evaluated using reference genomes from the same sample as the assemblies. Specifically, HG002 was evaluated against the HG002-T2T reference genome, while the other human datasets were assessed using high-quality assemblies published by the HPRC. For the assembly quality assessment on rice datasets (*48*), the reference genome used was the Nipponbare T2T genome. For the assembly quality assessment on Arabidopsis datasets (*49*), a high-quality Arabidopsis thaliana genome (*50*) was used as the reference.

Commands used by germline SVs detection and evaluation

Alignment-based SV detection and evaluation:

```
> minimap2 -ax map-hifi GRCH37.fa <input_file.fasta> -t 20 > aln.sam
> samtools view -Sb aln.sam > aln.bam
> samtools sort aln.bam -o aln_sorted.bam
> samtools index aln_sorted.bam
```

```
> svim alignment my_sample aln_sorted.bam GRCH37.fa
```

The SV detection algorithm based on contigs alignment was performed using svim-asm, with GRCh37 as the reference genome. The resulting files were sorted and indexed using samtools, and the “vcf” files were also sorted and indexed. The commands are as follows:

```
> minimap2 -a -x asm5 --cs -r2k -t 20 GRCH37.fa <your_assembly.fasta> > aln.sam
> samtools view -Sb aln.sam > aln.bam
> samtools sort aln.bam -o aln_sorted.bam
> samtools index aln_sorted.bam
> svim-asm haploid <your_dir> aln_sorted.bam GRCH37.fa
> bcftools sort <my.vcf> -o <my_sorted.vcf>
> bgzip <my_sorted.vcf>
> tabix <my_sorted.vcf.gz>
```

The evaluation was conducted using Truvari (v3.2.0) with the GIAB version 0.6 SV benchmark (high-confidence regions). The Challenging Medical Relevant Genes (CMRG) SV panel was also utilized for the assessment. The commands are as follows:

```
> truvari bench --passonly -p 0 --sizefilt 50 --sizemin 50 --sizemax 1000000 --includebed HG002_SVs_Tier1_v0.6.bed -b HG002_SVs_Tier1_v0.6.vcf.gz -c <my_sorted.vcf.gz> -f hs37d5.fa -o <my_dir>
> truvari bench --passonly -p 0 --sizefilt 50 --sizemin 50 --sizemax 1000000 --includebed HG002_GRCh37_CMRG_SV_v1.00.bed -b HG002_GRCh37_CMRG_SV_v1.00.vcf.gz -c <my_sorted.vcf.gz> -f hs37d5.fa -o <my_dir>
```

### Commands used by scaffolding, evaluation, and synteny analysis

The scaffolding was performed using Ragtag, with CHM13-T2T (v2.0) selected as the reference genome. The commands are as follows:

```
> ragtag.py scaffold <reference.fa> <query.fa>
```

The evaluation was similarly conducted using BUSCO and QUAST to assess basic metrics such as contiguity, with CHM13 (v2.0) selected as the reference genome. Assembly accuracy was evaluated using reference genomes consistent with the sample source of the assembled reads. Specifically, the reference genome for HG002 was HG002-T2T, while for the other human datasets, the reference genomes were the scaffolded sequences from high-quality assemblies published by HPRC, processed using Ragtag.

Synteny analysis was performed using NGenomeSyn. The commands are as follows:

```
> GetTwoGenomeSyn.pl -InGenomeA <your_scaffolding.fasta> - InGenomeB <your_reference.fasta> -OutPrefix <your_dir>
```

### Commands for pangenome graph construction and bubble region consistency evaluation

The pangenome graph was constructed using minigraph. The shell script is as follows:

```
> CHM13_path="<CHM13.fasta>"
> ref=$(find . -type f -path "<GRCH38.fasta>")
```

> other_sample_paths=$(find . -type f -path "*/hifiasm/output_name.fasta" | sort)
> sample_paths="$CHM13_path $other_sample_paths"
> cmd=" minigraph -cxggs -t20 $ref $sample_paths > GRCH38_hifiasm_minigraph_name.gfa"
> eval $cmd

The bubble regions of the pangenome were identified using gfatools, generating a BED file:
> gfatools bubble graph.gfa > var.bed

Using the BED file generated from the pangenome graph constructed by minigraph from the HPRC release as the reference (real.bed), bubble regions in our pangenome graph with over 80% overlap with real.bed are considered true positives (TP), otherwise they are considered false positives (FP). The commands are as follows:
> bedtools intersect -a <our_graph.bed> -b real.bed -f 0.80 -u > tp.bed
> bedtools intersect -a <our_graph.bed> -b real.bed -f 0.80 -v > fp.bed
> bedtools intersect -a <real.bed> -b <our_graph.bed> -f 0.80 -v >fn.bed

The evaluation was conducted across different intervals using our provided evalu-ate_bed.py, after modifying the paths of the input files.

Commands for large, rare germline SVs detection based on pangenome graph and evaluation

First, the input sequence is aligned to the reference genome GRCh38 using miniamp2-2.28+, followed by alignment to the reference genome CHM13-T2T. Finally, the input is aligned using minigraph-0.21+. When the input data is HiFi data, the com-mands are as follows:
> minimap2 -cx map-hifi -s50 --ds GRCH38.fa <your_hifi_dataset.fasta> -t <threads> >hifi_grch38.paf
> minimap2 -cx map-hifi -s50 --ds CHM13-T2T.fa <your_hifi_dataset.fasta> -t <threads> >hifi_chm13.paf
> minigraph -cxlr <pangraph.gfa> <hifi_dataset.fasta> -t <threads> > hifi_pan.gaf

When the input data is nanopore data, the commands are as follows:
> minimap2 -cx map-ont GRCH38.fa <your_ont_dataset.fasta> -t <threads> > ont_grch38.paf
> minimap2 -cx map-ont CHM13-T2T.fa <ont_dataset.fasta> -t <threads> > ont_chm13.paf
> minigraph -cxlr <pangraph.gfa> <your_hifi_dataset.fasta> -t <threads> > ont_pan.gaf

Germline-specific SV detection was performed using minisv.
> minisv.js e -b data/hs38.cen-mask.bed hifi_grch38.paf hifi_chm13.paf hifi_pan.gaf | bash > hifi_sv.rsv
> cat hifi.rsv | sort -k1,1 -k2,2 -S4g | minisv.js merge - > hifi_sv.msv
> minisv.js genvcf hifi_sv.msv > hifi_sv.vcf

Similarly, ont_sv.vcf can be obtained. Evaluation was performed using Truvari, with the SV benchmark set consisting of the SVs identified by minisv from the alignments to two linear reference genomes and the pangenome graph released by HPRC (re-al.vcf). All VCF files were sorted, compressed, and indexed, followed by the evaluation.

Commands for detecting large cancer somatic SVs

There are two scenarios for cancer somatic variant detection using minisv: detection with only tumor samples and detection with tumor-normal pairs. For detection with only tumor samples, the process is the same as long, rare germline SV detection based on the pangenome graph. In the case of tumor-normal pairs, miniamp2 is used to perform reciprocal alignment between the tumor and normal datasets, followed by detection using minisv. The commands are as follows:

```
> minimap2 -x ava-ont <my_normal.fasta> <my_tumor.fasta> > normal_tumor.paf
> minisv.js e -n TUMOR -0b data/hs38.cen-mask.bed ont_grch38.paf ont_chm13.paf ont_pan.gaf
normal_tumor.paf  > cancer.rsv
> minisv.js extract -n Normal normal.paf  > normal.rsv
> cat cancer.rsv normal.rsv | sort -k1,1 -k2,2 -S4g | minisv.js merge - | grep TUMOR | grep -v
NORMAL > paired.msv
> minisv.js genvcf paired.msv > paired.vcf
```

The evaluation was conducted using the same parameters as mentioned above for Truvari, except that the SV benchmark set from COLO829 was selected for the evaluation. The commands used by Sniffles for detecting cancer somatic SVs are as follows:

```
> sniffles --input mapped_input.bam --vcf output.vcf --mosaic
```

Pangenome graph augmentation commands

The scripts for graph augmentation are as follows:

```
> ref=hprc-v1.0-minigraph-grch38.gfa
> other_sample_paths=$(find . -type f -path "*/hificcl_name.fasta" | sort)
> sample_paths="$other_sample_paths"
> cmd="minigraph -cxggs -t20 $ref $sample_paths > plus_minigraph.gfa"
> eval $cmd
```

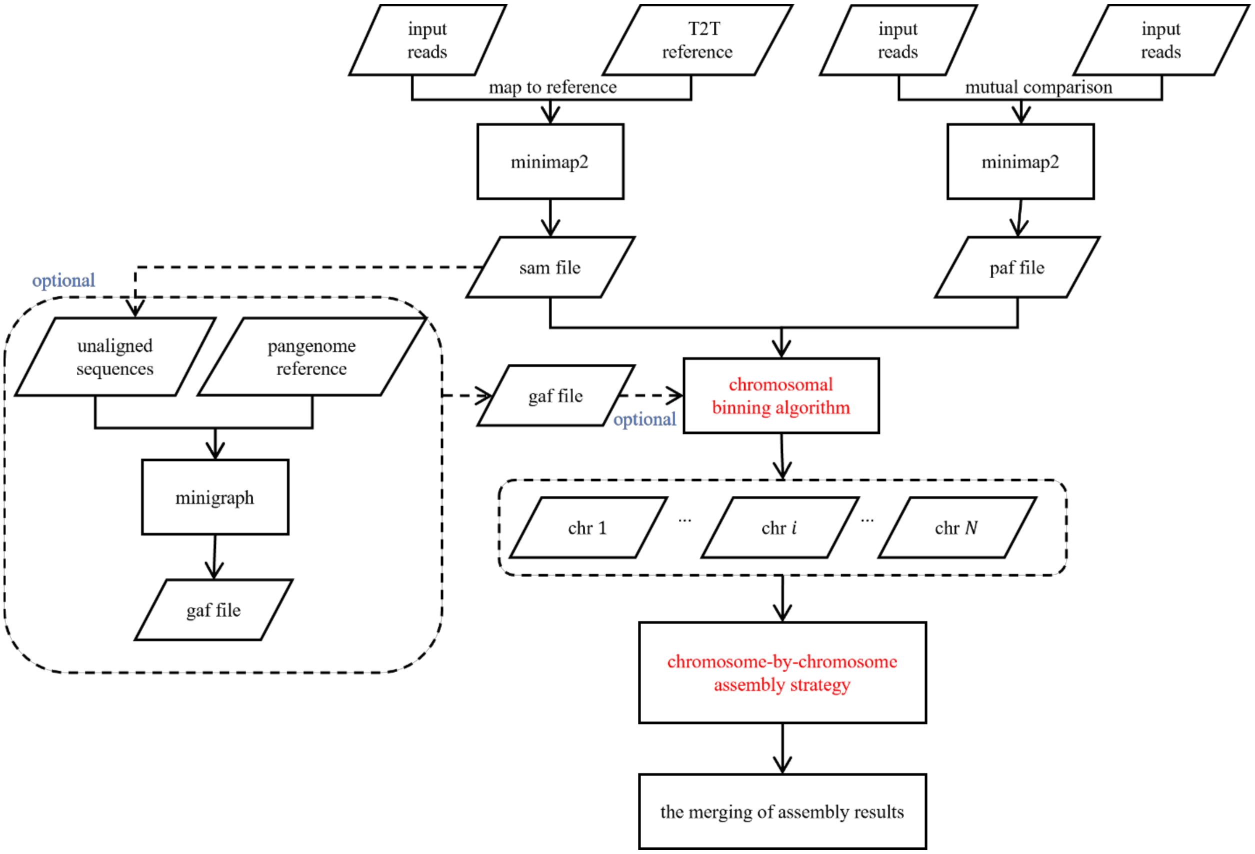


**Fig. S1. The whole pipeline of HiFiCCL.** HiFiCCL is divided into two kinds of modes, primary and optional modes. The primary mode operates by utilizing alignment information between reads and the linear reference, as well as the mutual comparison information among the reads themselves to guide the binning of reads by chromosome. The optional mode, building on the primary mode, further guides the binning process using alignment information of reads that did not align in the primary mode with pangenome graph. Finally, the assembly is completed by employing a chromosome-by-chromosome assembly strategy.

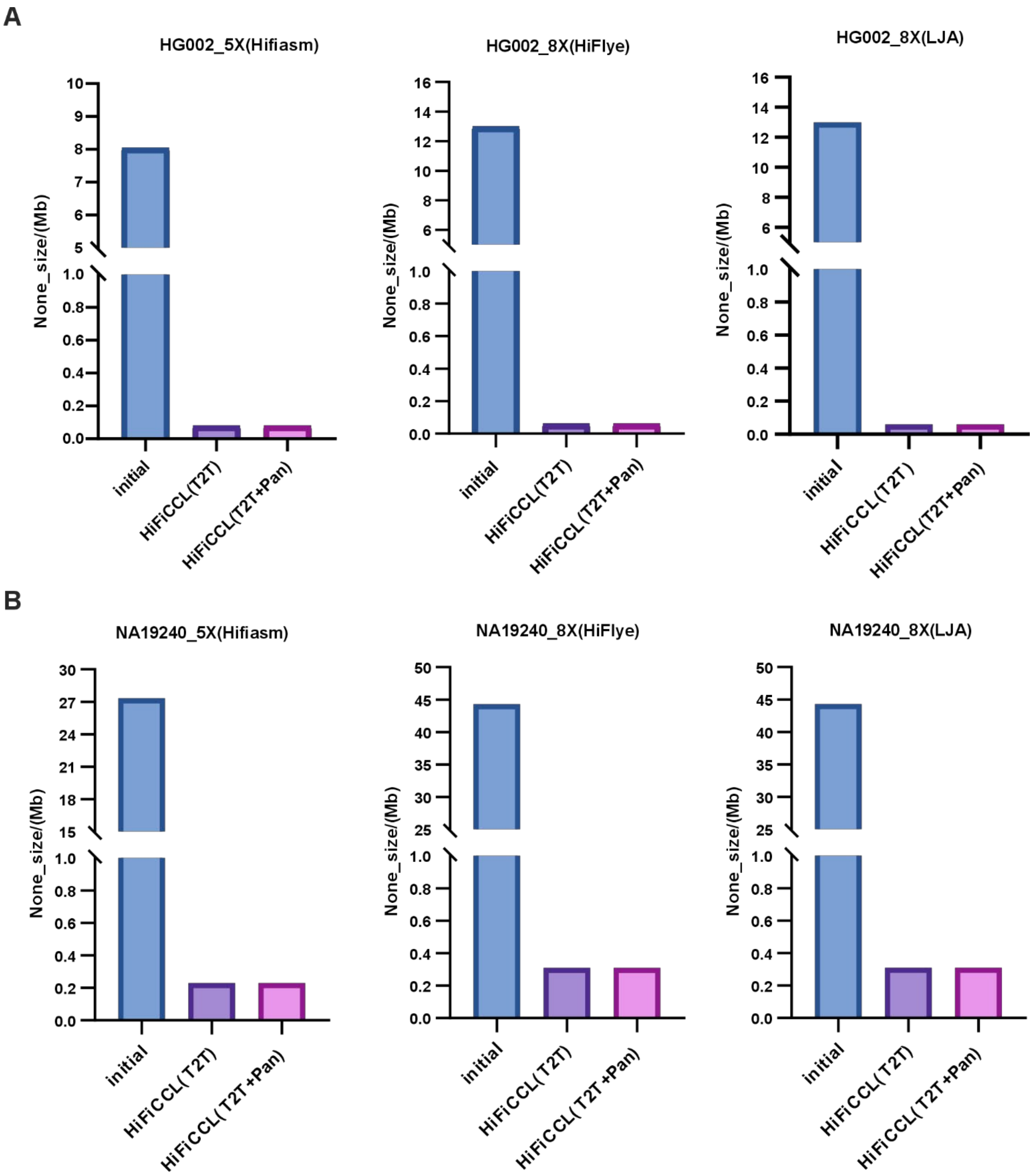


**Fig. S2. Comparison of the size of reads classified as 'None'.** (A) Comparison of the size of reads that did not align in the initial direct mapping versus after processing with the two modes of HiFiCCL for the HG002 dataset (5X, 8X). (B) Comparison of the size of reads that did not align in the initial direct mapping versus after processing with the two modes of HiFiCCL for the NA19240 dataset (5X, 8X).

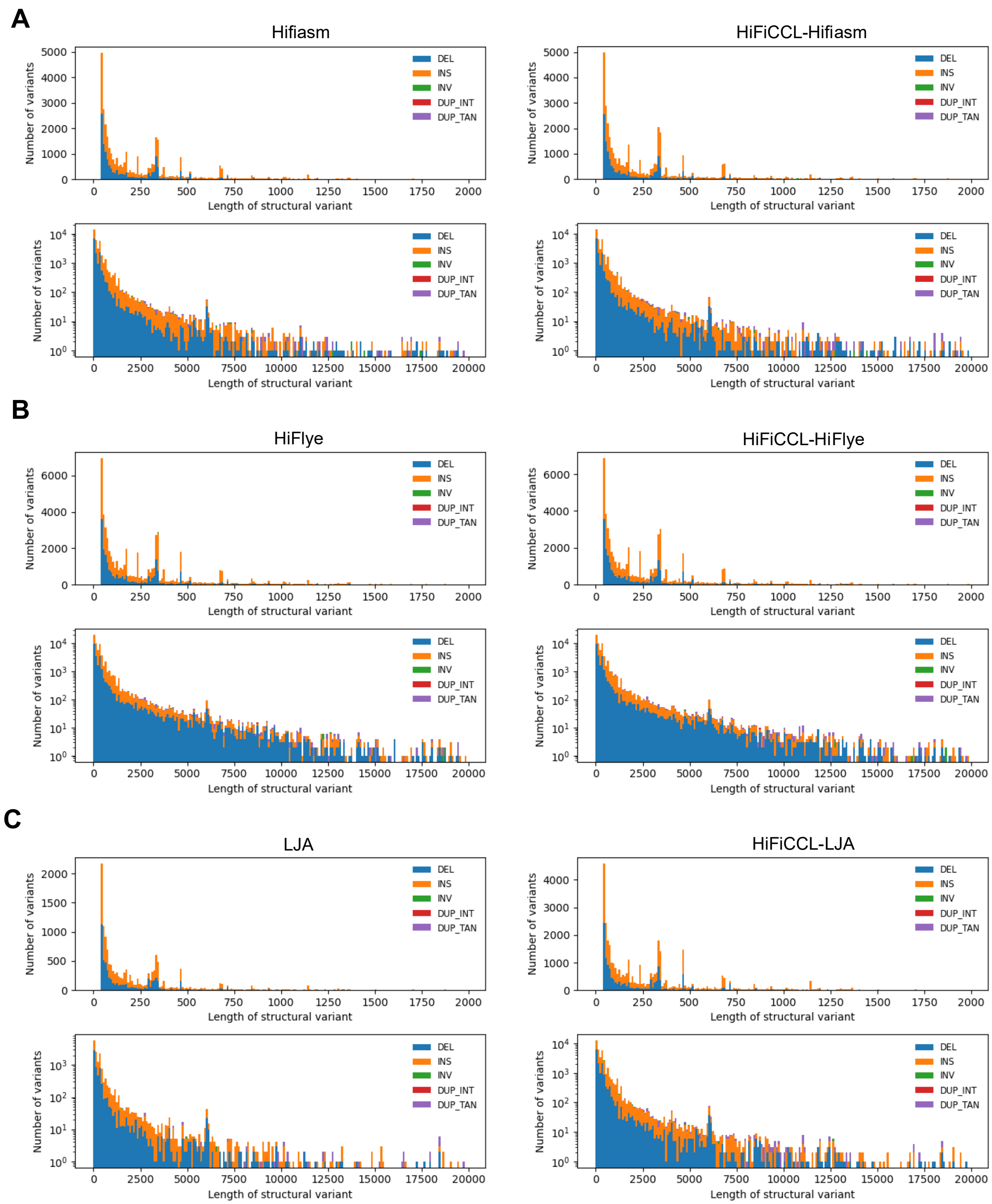


**Fig. S3. Statistical count of structural variations (SVs) of different sizes and types.** Statistical count of structural variations (SVs) of different sizes and types. (A) Statistical analysis of the count of different sizes and types of structural variations (SVs) detected based on the assembly results from Hifiasm and HiFiCCL-Hifiasm. (B) Statistics of different sizes and types of SVs identified through comparisons of the assembly results from HiFlye and HiFiCCL-HiFlye. (C) Count of different sizes and types of SVs detected based on the assembly results from LJA and HiFiCCL-LJA.

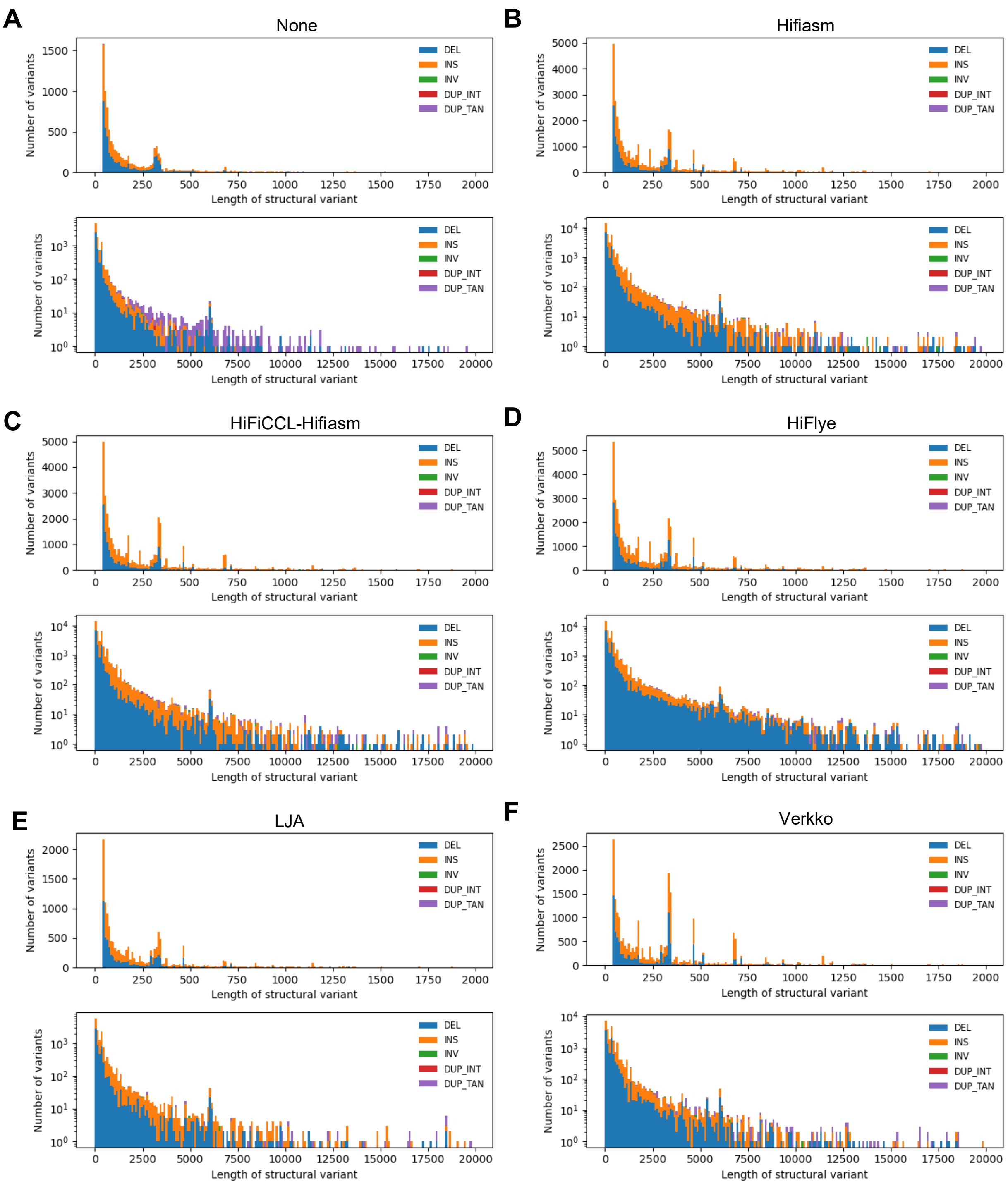


**Fig. S4. Statistical counts of structural variations (SVs) of different sizes and types.** Statistics and counts of structural variations (SVs) of different sizes and types were identified based on reads-alignment detection (A) and through alignment of assembly results from Hifiasm (B), HiFiCCL-Hifiasm (C), HiFlye (D), LJA (E), and Verkko (F).

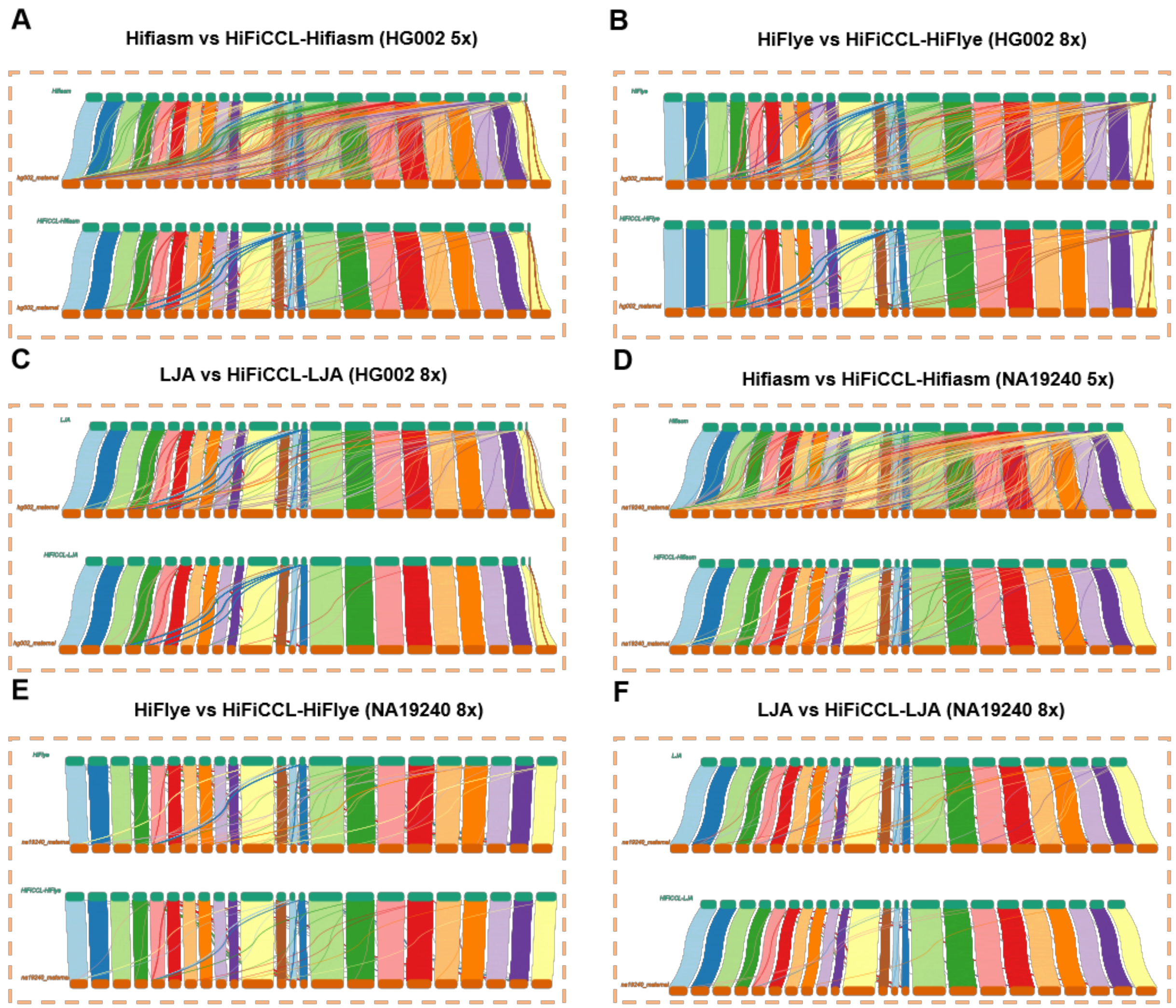


**Fig. S5. Synteny analysis of scaffolding results on ultra-low coverage HiFi datasets for HG002 and NA19240 (maternal).** In each subgraph, the top section shows the results based on the base assembler, while the bottom section displays the results after incorporating HiFiCCL. (A) Synteny analysis between the scaffolds from Hifiasm and HiFiCCL-Hifiasm assembly results post-scaffolding and the maternal reference genome HG002 at 5X coverage. (B) Synteny between the scaffolds from HiFlye and HiFiCCL-HiFlye assembly results post-scaffolding and the maternal reference genome HG002 at 8X coverage. (C) Synteny between the scaffolds from LJA and HiFiCCL-LJA assembly results post-scaffolding and the maternal reference genome HG002 at 8X coverage. (D) Synteny analysis between the scaffolds from Hifiasm and HiFiCCL-Hifiasm assembly results post-scaffolding and the maternal reference genome NA19240 at 5X coverage. (E) Synteny analysis between the scaffolds from HiFlye and HiFiCCL-HiFlye assembly results post-scaffolding and the maternal reference genome NA19240 at 8X coverage; f, synteny between the scaffolds from LJA and HiFiCCL-LJA assembly results post-scaffolding and the maternal reference genome NA19240 at 8X coverage.

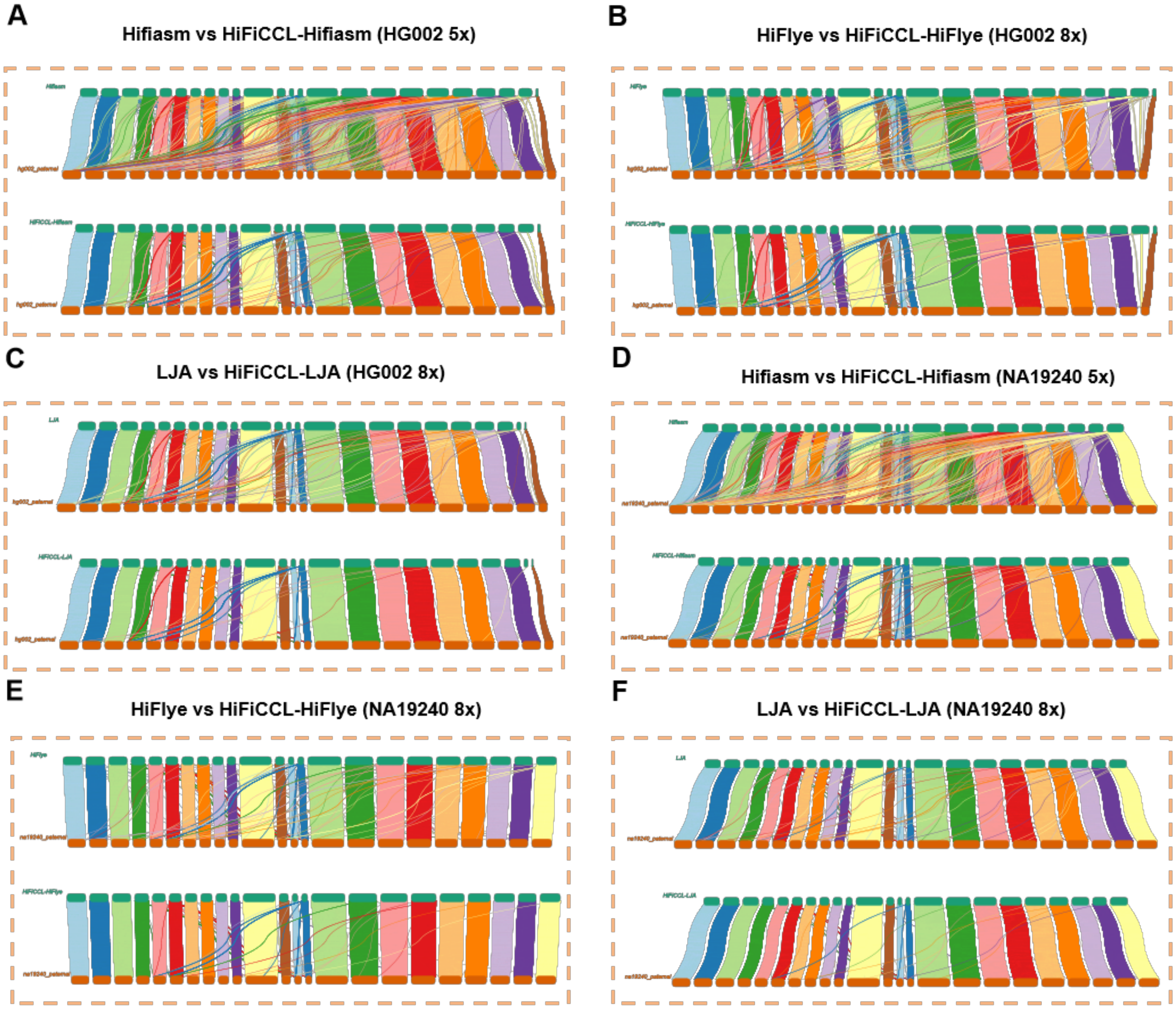


**Fig. S6. Synteny analysis of scaffolding results on ultra-low coverage HiFi datasets for HG002 and NA19240 (paternal).** In each subgraph, the top section shows the results based on the base assembler, while the bottom section displays the results after incorporating HiFiCCL. (A) Synteny analysis between the scaffolds from Hifiasm and HiFiCCL-Hifiasm assembly results post-scaffolding and the paternal reference genome HG002 at 5X coverage. (B) Synteny analysis between the scaffolds from HiFlye and HiFiCCL-HiFlye assembly results post-scaffolding and the paternal reference genome HG002 at 8X coverage. (C) Synteny between the scaffolds from LJA and HiFiCCL-LJA assembly results post-scaffolding and the paternal reference genome HG002 at 8X coverage. (D) Synteny between the scaffolds from Hifiasm and HiFiCCL-Hifiasm assembly results post-scaffolding and the paternal reference genome NA19240 at 5X coverage. (E) Synteny between the scaffolds from HiFlye and HiFiCCL-HiFlye assembly results post-scaffolding and the paternal reference genome NA19240 at 8X coverage. (F) Synteny between the scaffolds from LJA and HiFiCCL-LJA assembly results post-scaffolding and the paternal reference genome NA19240 at 8X coverage.

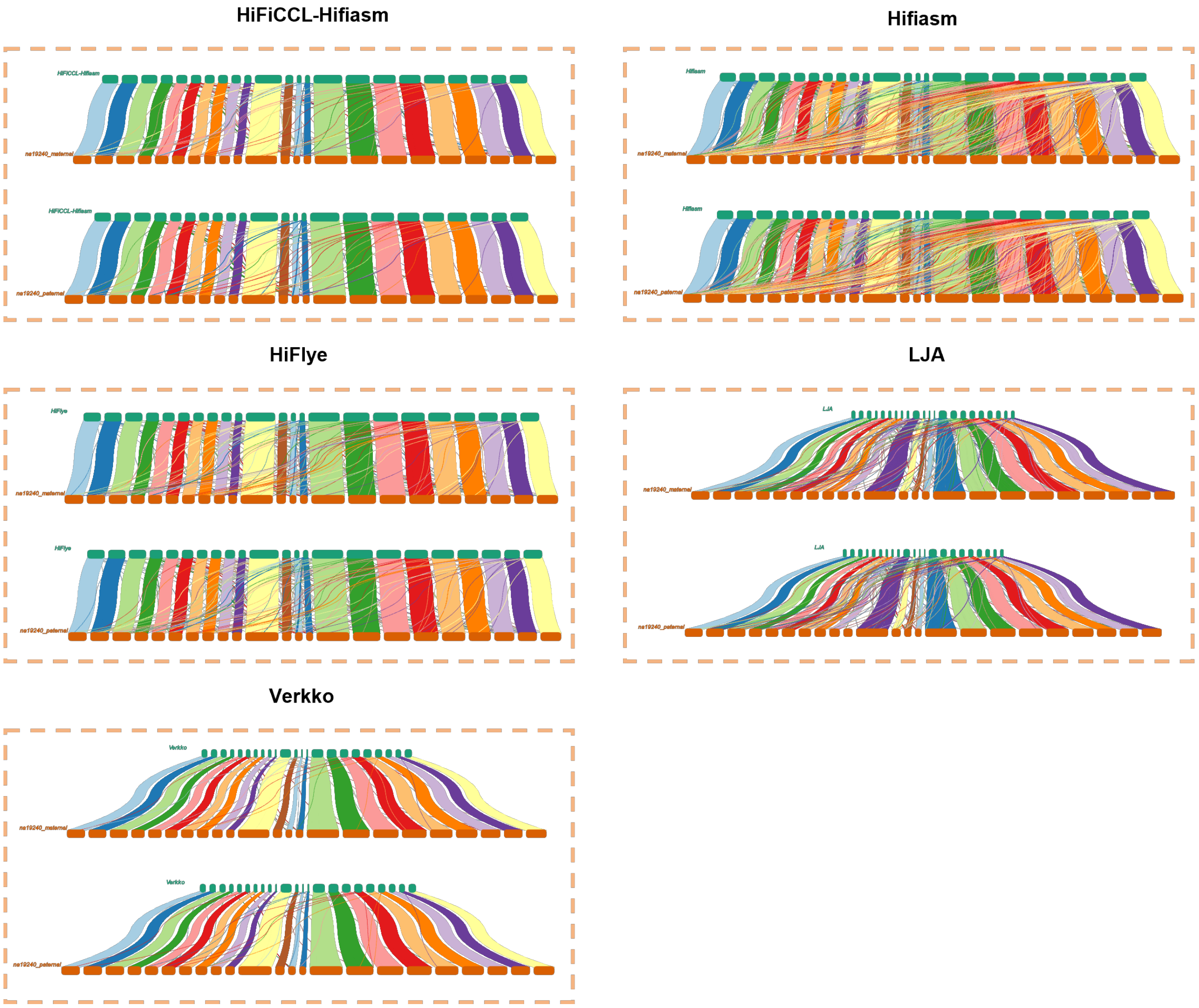


**Fig. S7. Synteny analysis of scaffolding results on ultra-low coverage HiFi dataset for NA19240.** It shows the synteny between the scaffolding performance of different assembler results and the maternal and paternal reference genomes for NA19240.

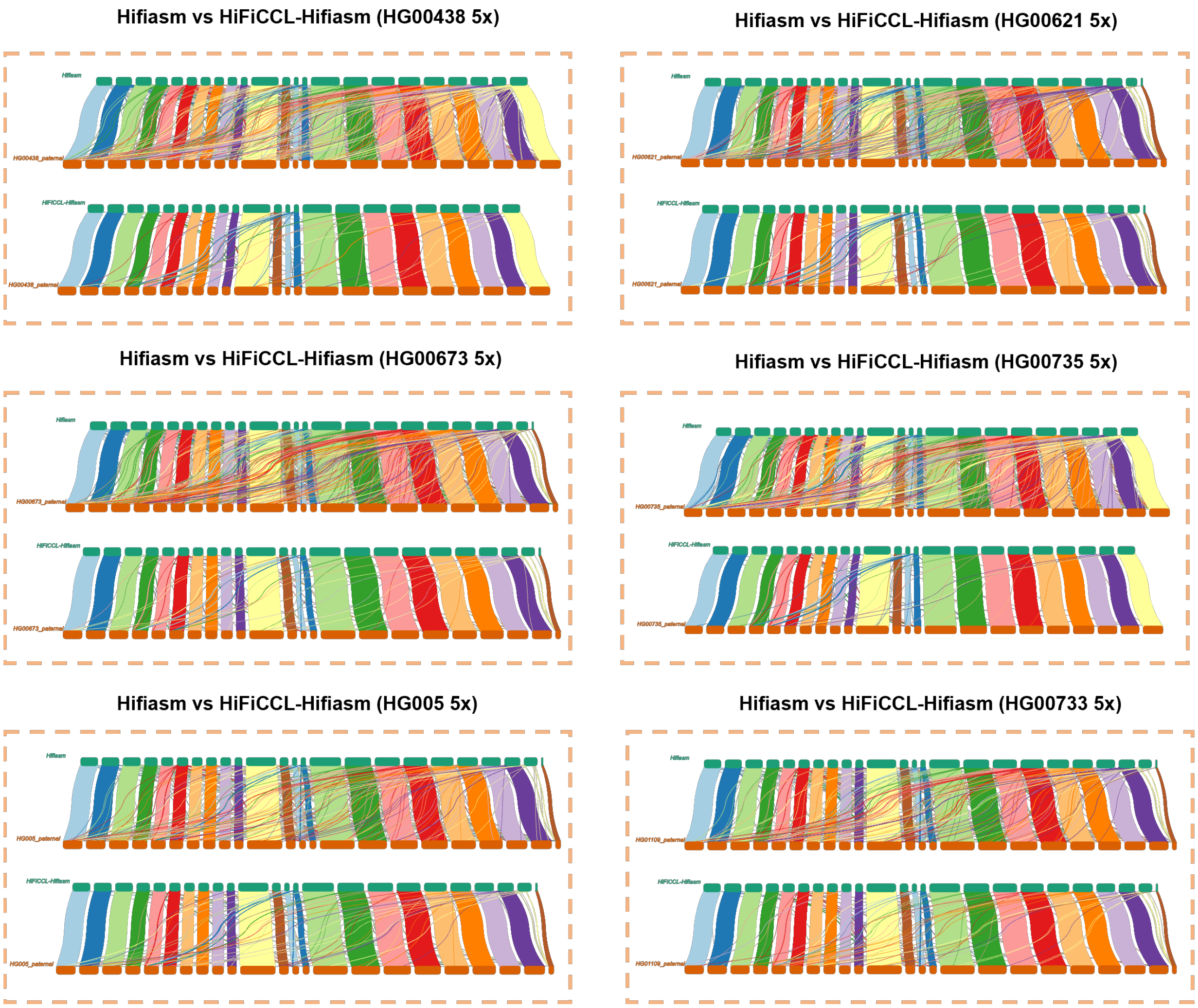


**Fig. S8. Synteny analysis of scaffolding results on multiple datasets.** In each box, the top section shows the results based on the base assembler, while the bottom section displays the results after incorporating HiFiCCL. It shows the scaffolding performance of HiFiCCL-Hifiasm and Hifiasm assemblies, with the paternal reference genome used as the reference.

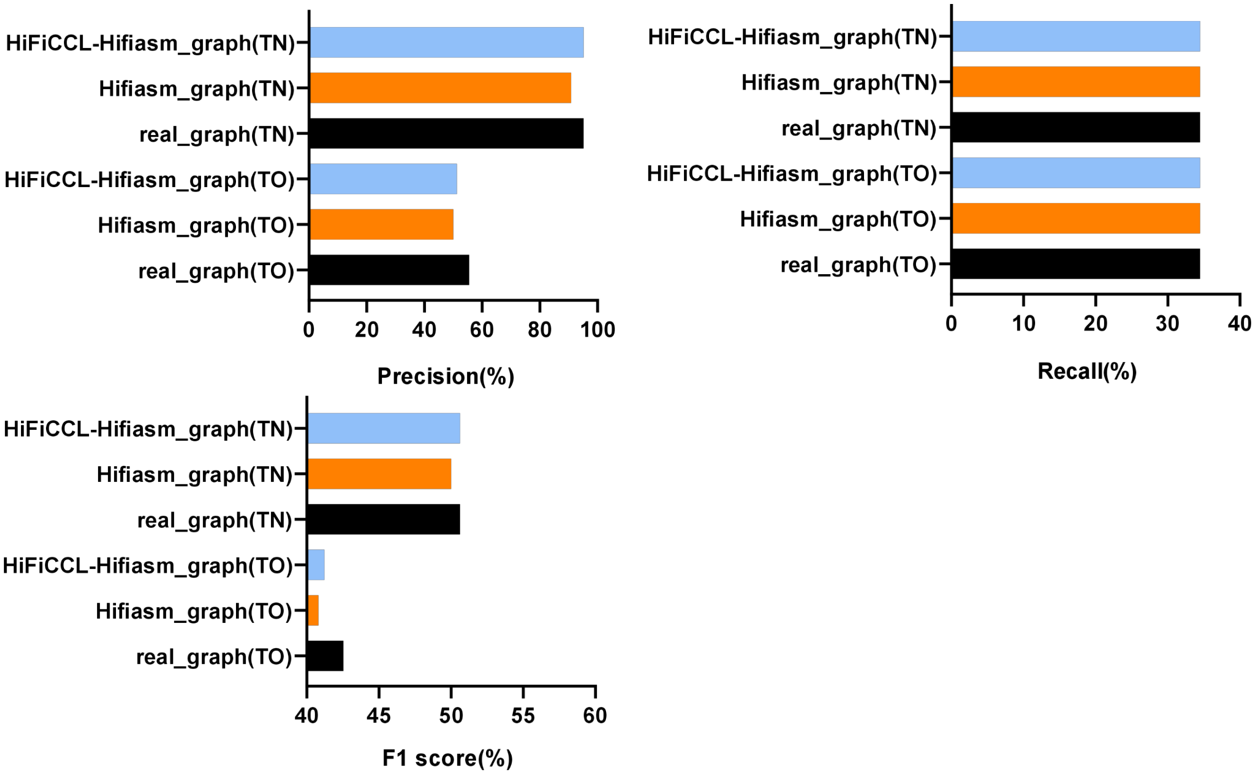


**Fig. S9. Tumor somatic SVs detection using two modes of minisv.** It displays the detection of large somatic SVs in the COLO829 dataset using the pan-genome graph released by the HPRC, and those constructed from Hifiasm and HiFiCCL-Hifiasm assembly results, with TO indicating the use of only tumor tissue datasets, and TN indicating the use of both tumor cell and normal cell datasets

**Table S1. Statistics of human primary assemblies across different assemblers.**

| Dataset | Assembler | Size (Gb) | Contigs number | MCL (Mb) | NG50 (Kb) | NGA50 (Kb) | Gene completeness (BUSCO) | |
|---|---|---|---|---|---|---|---|---|
| | | | | | | | **Complete /Single(%)** | **Missing (%)** |
| HG002 (HiFi 5x) | HiFiCCL-Hifiasm | 2.79 | 18304 | 274.16 | 199.05 | 185.04 | 82.6/79.1 | 7.1 |
| | Hifiasm | 2.73 | 17868 | 331.84 | 194.55 | 174.55 | 82.0/78.7 | 7.6 |
| | HiFlye | 3.35 | 39567 | 501.71 | 199.45 | 175.02 | 84.3/76.9 | 6.1 |
| | LJA | 1.7 | 11913 | 211.1 | 83.1 | 70.5 | 47.6/45.2 | 43.0 |
| | Verkko | 2.0 | 43952 | 49.0 | 31.4 | 29.5 | 42.3/36.9 | 44.8 |
| | GALA | - | - | - | - | - | - | - |
| NA19240 (HiFi 5x) | HiFiCCL-Hifiasm | 2.64 | 23417 | 179.15 | 124.83 | 116.19 | 77.9/74.9 | 9.9 |
| | Hifiasm | 2.57 | 22561 | 245.96 | 123.48 | 112.86 | 77.5/74.5 | 10.1 |
| | HiFlye | 3.06 | 43236 | 259.01 | 134.33 | 125.70 | 77.4/70.8 | 10.3 |
| | LJA | 0.64 | 4955 | 115.58 | - | - | 22.2/21.8 | 72.1 |
| | Verkko | 1.06 | 24974 | 27.42 | - | - | 26.7/23.2 | 63.1 |
| | GALA | - | - | - | - | - | - | - |

**Table S2. Comparison of runtime and memory usage on human datasets.**

| Dataset | Assembler | Elapsed (wall clock) time /(h:mm:ss) | Maximum resident set size /(kbytes) |
|---|---|---|---|
| HG002 (5x) | HiFiCCL-Hifiasm | 5:02:09 | 55,754,616 |
| | Hifiasm | 9:08:55 | 51,458,516 |
| | HiFlye | 4:29:27 | 48,036,364 |
| | LJA | 8:04:22 | 74,394,632 |
| | Verkko | 5:27:08 | 19,012,472 |
| | GALA | - | - |
| NA19240 (5x) | HiFiCCL-Hifiasm | 4:09:50 | 53,132,656 |
| | Hifiasm | 6:32:53 | 51,499,152 |
| | HiFlye | 3:43:48 | 47,933,488 |
| | LJA | 6:24:33 | 77,701,724 |
| | Verkko | 4:11:51 | 18,995,300 |
| | GALA | - | - |

**Table S3. Comparison of chromosomal clustering time and memory usage between HiFiCCL and GALA on human datasets.**

| Dataset | Assembler | Elapsed (wall clock) time /(h:mm:ss) | Maximum resident set size /(kbytes) |
|---|---|---|---|
| HG002 (5x) | HiFiCCL-clustering | 03:17:16 | 54,219,140 |
| | GALA-clustering | 62:13:07 | 52,018,932 |
| NA19240 (5x) | HiFiCCL-clustering | 02:44:10 | 53,795,588 |
| | GALA-clustering | 56:39:51 | 44,448,032 |

**Table S4. Statistics of plant primary assemblies across different assemblers.**

| Dataset | Assembler | Size (Mb) | Contigs number | MCL (Mb) | NG50 (Kb) | NGA50 (Kb) | Gene completeness (BUSCO) | |
|---|---|---|---|---|---|---|---|---|
| | | | | | | | Complete /Single(%) | Missing (%) |
| Rice (HiFi 5x) | HiFiCCL-Hifiasm | 321.8 | 5362 | 38.0 | 61.0 | 57.4 | 85.4/83.2 | 11.4 |
| | Hifiasm | 297.8 | 3760 | 74.3 | 75.1 | 64.3 | 78.6/76.7 | 17.9 |
| | LJA | 12.9 | 258 | 2.5 | - | - | 3.5/3.5 | 96.1 |
| | verkko | 83.4 | 846 | 4.2 | - | - | 25.6/25.1 | 73.5 |
| | GALA | - | - | - | - | - | - | - |
| Arabidopsist haliana (HiFi 5x) | HiFiCCL-Hifiasm | 117.6 | 1380 | 13.3 | 90.4 | 86.9 | 87.6/86.6 | 10.0 |
| | Hifiasm | 115.9 | 1340 | 15.6 | 90.5 | 85.9 | 87.1/86.2 | 10.5 |
| | LJA | - | - | - | - | - | - | - |
| | verkko | 45.2 | 495 | 6.4 | - | - | 27.4/26.8 | 71.6 |
| | GALA | - | - | - | - | - | - | - |

**Table S5. Comparison of runtime and memory usage on plant datasets.**

| Dataset | Assembler | Elapsed (wall clock) time /(h:mm:ss) | Maximum resident set size /(kbytes) |
|---|---|---|---|
| Rice (HiFi 5x) | HiFiCCL-Hifiasm | 00:40:34 | 19,021,336 |
| | Hifiasm | 00:20:02 | 21,519,320 |
| | LJA | 01:05:44 | 9,723,420 |
| | Verkko | 00:21:08 | 18,876,364 |
| | GALA | - | - |
| Arabidopsisthaliana (HiFi 5x) | HiFiCCL-Hifiasm | 00:12:02 | 17,425,656 |
| | Hifiasm | 00:05:01 | 18,261,228 |
| | LJA | 00:09:00 | 3,485,796 |
| | Verkko | 00:14:14 | 18,391,424 |
| | GALA | - | - |

**Table S6. Comparison of chromosomal clustering time and memory usage between HiFiCCL and GALA on plant datasets.**

| Dataset | Assembler | Elapsed (wall clock) time /(h:mm:ss) | Maximum resident set size /(kbytes) |
|---|---|---|---|
| Rice (HiFi 5x) | HiFiCCL-clustering | 00:26:26 | 19,736,972 |
| | GALA-clustering | 04:23:50 | 20,594,464 |
| Arabidopsi sthaliana (HiFi 5x) | HiFiCCL-clustering | 00:07:31 | 8,241,860 |
| | GALA-clustering | 01:05:25 | 3,322,192 |

**Table S7. Statistics of human primary assemblies on the 45 human datasets (~5x).** The bold data indicates that the HiFiCCL metric's performance surpassed that of the base assembler.

| Dataset | Assembler | Size (Gb) | Contigs number | MCL (Mb) | NG50 (Kb) | NGA50 (Kb) | Gene completeness (BUSCO) | |
|---|---|---|---|---|---|---|---|---|
| | | | | | | | Complete /Single(%) | Missing (%) |
| HG00438 | HiFiCCL-Hifiasm | **2.63** | 18573 | **170.77** | 155.91 | 143.67 | **77.1/74.3** | **11.5** |
| | Hifiasm | 2.58 | 17636 | 254.39 | 159.07 | 143.68 | 76.5/73.7 | 12.1 |
| HG00621 | HiFiCCL-Hifiasm | **2.62** | 19071 | **172.33** | 150.56 | 140.15 | **77.5/74.8** | **10.6** |
| | Hifiasm | 2.57 | 17892 | 235.82 | 156.74 | 143.15 | 77.1/74.2 | 11.1 |
| HG00673 | HiFiCCL-Hifiasm | **2.65** | 19409 | **195.75** | **153.54** | **143.07** | **79.8/77.0** | **8.8** |
| | Hifiasm | 2.57 | 18607 | 264.49 | 150.87 | 137.22 | 79.4/76.6 | 9.3 |
| HG00735 | HiFiCCL-Hifiasm | **2.66** | 18917 | **181.84** | **157.22** | **146.16** | **75.7/72.9** | **12.3** |
| | Hifiasm | 2.59 | 18398 | 250.60 | 154.65 | 140.50 | 74.9/72.4 | 12.8 |
| HG00741 | HiFiCCL-Hifiasm | **2.65** | 17434 | **188.86** | **168.79** | **156.14** | **76.5/73.5** | **12.5** |
| | Hifiasm | 2.59 | 17007 | 248.75 | 163.40 | 149.09 | 75.4/72.6 | 13.6 |
| HG005 | HiFiCCL-Hifiasm | **2.69** | 20947 | 233.20 | 150.82 | 140.48 | 78.9/76.3 | 9.1 |
| | Hifiasm | 2.66 | 20475 | 218.63 | 152.20 | 140.71 | 78.9/76.3 | 9.1 |
| HG01109 | HiFiCCL-Hifiasm | **2.66** | 20199 | **207.84** | 148.04 | 136.71 | 79.8/76.8 | **9.3** |
| | Hifiasm | 2.65 | 19782 | 208.81 | 150.43 | 137.72 | 80.1/77.1 | 9.4 |
| HG01123 | HiFiCCL-Hifiasm | **2.69** | 23569 | **161.93** | **132.59** | **125.41** | **78.2/75.1** | **9.9** |
| | Hifiasm | 2.62 | 22901 | 263.63 | 128.85 | 119.24 | 77.2/74.4 | 10.5 |
| HG01175 | HiFiCCL-Hifiasm | **2.67** | 19451 | **178.63** | 154.43 | 143.36 | 79.2/76.4 | **10.0** |
| | Hifiasm | 2.66 | 18951 | 190.90 | 157.58 | 145.51 | 79.2/76.5 | 10.1 |
| HG01106 | HiFiCCL-Hifiasm | **2.67** | 18653 | 206.41 | **165.14** | 150.32 | **79.3/76.7** | **9.7** |
| | Hifiasm | 2.66 | 18319 | 199.11 | 165.10 | 151.09 | 79.2/76.5 | 9.8 |
| HG01243 | HiFiCCL-Hifiasm | **2.67** | 20978 | **213.91** | **144.72** | **133.33** | **79.6/76.9** | **8.8** |
| | Hifiasm | 2.60 | 20312 | 265.49 | 140.78 | 127.93 | 78.8/76.2 | 9.6 |
| HG01258 | HiFiCCL-Hifiasm | **2.62** | 21108 | **220.54** | **135.51** | **126.43** | **74.3/71.6** | **12.1** |
| | Hifiasm | 2.55 | 20182 | 259.35 | 134.56 | 123.30 | 73.6/71.0 | 12.7 |
| HG01071 | HiFiCCL-Hifiasm | **2.64** | 15953 | 200.38 | 183.15 | 170.57 | 78.5/75.9 | **11.7** |
| | Hifiasm | 2.63 | 15640 | 197.20 | 186.67 | 172.48 | 78.5/75.9 | 11.8 |
| HG01358 | HiFiCCL-Hifiasm | **2.68** | 22168 | **221.66** | **137.07** | **127.96** | **78.3/75.6** | **8.9** |
| | Hifiasm | 2.60 | 21303 | 281.14 | 134.13 | 122.62 | 77.1/74.5 | 9.8 |
| HG01361 | HiFiCCL-Hifiasm | **2.67** | 20495 | **225.78** | **148.69** | **139.05** | **79.5/76.3** | **8.9** |
| | Hifiasm | 2.60 | 19830 | 287.91 | 145.15 | 132.34 | 78.3/75.2 | 9.8 |
| HG01891 | HiFiCCL-Hifiasm | **2.82** | 19157 | **225.60** | **193.65** | **176.07** | **84.6/81.3** | **6.2** |
| | Hifiasm | 2.78 | 19091 | 335.18 | 186.14 | 166.41 | 84.0/80.7 | 6.6 |
| HG01928 | HiFiCCL-Hifiasm | **2.65** | 17093 | **194.94** | **175.16** | **162.37** | **78.5/76.0** | **11.1** |
| | Hifiasm | 2.57 | 16546 | 263.86 | 170.02 | 154.17 | 77.6/75.2 | 11.8 |
| HG02080 | HiFiCCL-Hifiasm | **2.69** | 19363 | **187.28** | 158.06 | 145.70 | 78.8/76.1 | 9.5 |
| | Hifiasm | 2.67 | 18863 | 192.61 | 160.53 | 147.59 | 79.2/76.5 | 9.4 |
| HG00733 | HiFiCCL-Hifiasm | **2.60** | 28231 | 226.84 | 96.63 | **91.51** | **73.2/70.7** | **11.9** |
| | Hifiasm | 2.56 | 27106 | 218.75 | 97.21 | 91.28 | 71.4/69.0 | 13.1 |
| HG02109 | HiFiCCL-Hifiasm | **2.66** | 21222 | **167.54** | **140.04** | **129.96** | **78.7/75.8** | **9.4** |
| | Hifiasm | 2.59 | 20539 | 240.29 | 137.81 | 125.09 | 78.1/75.3 | 9.9 |
| HG02145 | HiFiCCL-Hifiasm | **2.61** | 21147 | **212.90** | 132.77 | 120.23 | 80.7/**77.5** | 8.0 |
| | Hifiasm | 2.60 | 20788 | 215.17 | 134.41 | 120.57 | 80.7/77.4 | 7.9 |
| HG02148 | HiFiCCL-Hifiasm | **2.67** | 21747 | **204.62** | **142.61** | **133.12** | **79.9/77.0** | **8.4** |

| | | | | | | | | |
|---|---|---|---|---|---|---|---|---|
| | Hifiasm | 2.59 | 20836 | 266.78 | 139.94 | 128.14 | 78.5/75.8 | 9.4 |
| HG01952 | HiFiCCL-Hifiasm | **2.66** | 18871 | 193.88 | 159.98 | 148.31 | 79.3/76.1 | 9.2 |
| | Hifiasm | 2.65 | 18471 | 182.26 | 162.38 | 150.44 | 79.7/76.5 | 9.0 |
| HG01978 | HiFiCCL-Hifiasm | **2.65** | 16529 | 208.66 | 177.32 | 164.02 | **77.9/74.7** | **11.1** |
| | Hifiasm | 2.64 | 16172 | 198.16 | 180.67 | 166.76 | 77.6/74.4 | 11.4 |
| HG02257 | HiFiCCL-Hifiasm | **2.69** | 21801 | **188.96** | 140.73 | 130.21 | 78.6/75.6 | 9.1 |
| | Hifiasm | 2.68 | 21360 | 191.95 | 142.87 | 131.38 | 78.6/75.7 | 8.9 |
| HG02486 | HiFiCCL-Hifiasm | **2.68** | 22999 | **178.65** | **132.72** | **122.48** | **77.1/73.7** | **9.9** |
| | Hifiasm | 2.60 | 22307 | 264.28 | 129.15 | 116.98 | 76.1/72.9 | 10.4 |
| HG02559 | HiFiCCL-Hifiasm | **2.67** | 20959 | **168.24** | 143.79 | 132.23 | 75.8/73.1 | 11.1 |
| | Hifiasm | 2.66 | 20572 | 177.69 | 146.06 | 133.57 | 76.0/73.3 | 10.9 |
| HG02572 | HiFiCCL-Hifiasm | **2.63** | 22755 | **220.85** | **125.03** | **112.07** | **76.2/73.3** | **12.0** |
| | Hifiasm | 2.57 | 22186 | 272.54 | 122.07 | 108.23 | 75.5/72.6 | 12.6 |
| HG02622 | HiFiCCL-Hifiasm | **2.67** | 18786 | 217.53 | 158.41 | 145.32 | 79.6/76.7 | **9.3** |
| | Hifiasm | 2.66 | 18402 | 211.35 | 161.16 | 147.29 | 79.7/76.8 | 9.4 |
| HG02630 | HiFiCCL-Hifiasm | **2.67** | 22264 | **209.98** | **135.06** | **125.10** | **77.8/74.8** | **9.3** |
| | Hifiasm | 2.60 | 21546 | 257.56 | 131.90 | 120.08 | 76.9/73.7 | 10.0 |
| HG02717 | HiFiCCL-Hifiasm | **2.79** | 20037 | **264.71** | 179.06 | 162.91 | **84.2/81.1** | **6.3** |
| | Hifiasm | 2.75 | 18898 | 345.19 | 184.08 | 163.92 | 83.4/80.4 | 7.0 |
| HG02723 | HiFiCCL-Hifiasm | **2.68** | 25284 | 200.93 | 121.85 | 113.71 | 79.7/76.5 | 8.0 |
| | Hifiasm | 2.67 | 24517 | 200.13 | 124.31 | 115.64 | 80.1/77.1 | 7.8 |
| HG02818 | HiFiCCL-Hifiasm | **2.69** | 24493 | **250.31** | **124.48** | **116.38** | **78.0/75.1** | **8.7** |
| | Hifiasm | 2.61 | 23854 | 279.20 | 121.15 | 111.02 | 76.8/74.1 | 9.6 |
| HG02886 | HiFiCCL-Hifiasm | **2.67** | 23437 | **184.45** | 129.53 | 120.65 | **76.7**/73.5 | 10.4 |
| | Hifiasm | 2.66 | 22900 | 188.71 | 131.65 | 122.40 | 76.6/73.6 | 10.4 |
| HG03453 | HiFiCCL-Hifiasm | **2.68** | 21135 | 201.41 | 143.15 | 131.87 | 79.8/76.7 | 8.6 |
| | Hifiasm | 2.67 | 20692 | 197.39 | 145.91 | 133.94 | 80.0/76.9 | 8.5 |
| HG03486 | HiFiCCL-Hifiasm | **2.65** | 22285 | **206.87** | **132.43** | **122.00** | **79.1/76.1** | **8.9** |
| | Hifiasm | 2.59 | 21398 | 255.42 | 132.16 | 119.23 | 78.6/75.5 | 9.6 |
| HG03492 | HiFiCCL-Hifiasm | **2.65** | 20034 | **215.16** | **148.89** | **136.81** | **79.6/76.7** | **9.7** |
| | Hifiasm | 2.57 | 19265 | 272.54 | 145.29 | 131.21 | 79.0/76.1 | 10.1 |
| HG03516 | HiFiCCL-Hifiasm | 2.67 | 23906 | **173.34** | 128.77 | 118.77 | 76.6/73.2 | **10.8** |
| | Hifiasm | 2.67 | 22311 | 185.85 | 134.77 | 123.60 | 76.8/73.4 | 10.9 |
| HG03540 | HiFiCCL-Hifiasm | 2.66 | 20746 | **202.87** | 142.98 | 131.83 | 78.5/75.6 | 9.4 |
| | Hifiasm | 2.66 | 20377 | 209.93 | 145.61 | 133.25 | 78.6/75.9 | 9.2 |
| HG03579 | HiFiCCL-Hifiasm | **2.64** | 21667 | **208.55** | 134.18 | 121.89 | 81.0/77.4 | **7.7** |
| | Hifiasm | 2.63 | 21173 | 211.46 | 136.98 | 124.07 | 81.1/77.4 | 7.8 |
| HG02055 | HiFiCCL-Hifiasm | **2.67** | 23499 | **175.22** | **127.77** | **117.86** | **74.6/71.7** | **11.7** |
| | Hifiasm | 2.60 | 22919 | 240.66 | 123.55 | 112.26 | 73.9/71.2 | 12.3 |
| HG03098 | HiFiCCL-Hifiasm | **2.67** | 21941 | **187.67** | **137.26** | **126.09** | **77.3/74.5** | **10.5** |
| | Hifiasm | 2.60 | 21332 | 254.50 | 134.09 | 121.13 | 76.5/74.0 | 11.2 |
| NA18906 | HiFiCCL-Hifiasm | 2.54 | 20288 | **191.55** | 128.55 | 117.74 | 78.1/75.2 | 10.3 |
| | Hifiasm | 2.54 | 18580 | 247.60 | 144.00 | 129.59 | 78.2/75.6 | 10.0 |
| NA21309 | HiFiCCL-Hifiasm | 2.65 | 24889 | **169.35** | 119.38 | 111.97 | 75.6/73.0 | 10.4 |
| | Hifiasm | 2.65 | 24010 | 175.45 | 124.29 | 116.15 | 76.0/73.5 | 10.3 |
| NA20129 | HiFiCCL-Hifiasm | **2.64** | 23529 | **309.73** | 124.02 | **116.01** | **75.9/72.9** | **10.4** |
| | Hifiasm | 2.60 | 22564 | 373.13 | 125.42 | 115.87 | 75.5/72.7 | 11.1 |

**Table S8. Statistics of human primary assemblies on the HG002 dataset at various coverage levels.** Bolded data indicates that the HiFiCCL metrics performance surpassed that of the base assembler, while a star in the top right corner denotes the best performance achieved.

| Dataset | Assembler | Size (Gb) | Contigs number | MCL (Mb) | NG50 (Kb) | NGA50 (Kb) | Gene completeness (busco) | |
|---|---|---|---|---|---|---|---|---|
| | | | | | | | Complete /Single(%) | Missing (%) |
| HG002 (HiFi 3x) | Hifiasm | 2.19 | 27581 | 192.59 | 65.66 | 62.20 | 61.5/59.1 | 22.0 |
| | HiFiCCL-Hifiasm | **2.30*** | 28881 | **180.93** | **69.74** | **66.28** | **63.1/60.5*** | **20.9** |
| | HiFiCCL-Hifiasm (optional) | **2.30*** | 28935 | **180.46*** | **69.78*** | **66.34*** | **63.2*/60.5*** | **20.8*** |
| | HiFlye | 2.13 | 32188 | 151.06 | 56.77* | 53.22* | 54.5/51.3 | 30.1 |
| | HiFiCCL-HiFlye | 2.12 | 32238 | **119.06** | 55.34 | 52.18 | **55.2*/52.2*** | **29.8*** |
| | HiFiCCL-HiFlye) (optional) | 2.12 | 32235 | **118.38*** | 55.39 | 52.27 | **55.2*/52.2*** | **29.8*** |
| | LJA | - | - | - | - | - | - | - |
| | HiFiCCL-LJA | - | - | - | - | - | - | - |
| | HiFiCCL-LJA (optional) | - | - | - | - | - | - | - |
| HG002 (HiFi 5x) | Hifiasm | 2.73 | 17868 | 331.84 | 194.55 | 174.55 | 82.0/78.7 | 7.6 |
| | HiFiCCL-Hifiasm | **2.79*** | 18304 | **274.16** | **199.05** | **185.04** | **82.6*/79.1*** | **7.1*** |
| | HiFiCCL-Hifiasm (optional) | **2.79*** | 18264 | **274.07*** | **198.76** | **184.97** | **82.6/*79.0** | **7.1*** |
| | HiFlye | 3.35 | 39567 | 501.71 | 199.45 | 175.02 | 84.3/76.9 | 6.1 |
| | HiFiCCL-HiFlye | **3.34** | **39564** | **422.82** | **200.25*** | **177.66** | **84.8/77.1** | 6.1 |
| | HiFiCCL-HiFlye (optional) | **3.34** | **39417*** | **416.26*** | 199.52 | **177.76*** | **85.1*/77.8*** | **5.8*** |
| | LJA | 1.76 | 11913 | 211.11 | 83.10 | 70.50 | 47.6/45.2 | 43.0 |
| | HiFiCCL-LJA | 1.73 | **10753*** | **100.10*** | 81.68 | **71.54** | 47.3/45.1 | 44.6 |
| | HiFiCCL-LJA (optional) | 1.73 | **10792** | **101.48** | 81.88 | **71.83*** | 47.2/45.0 | 44.6 |
| HG002 (HiFi 8x) | Hifiasm | 3.61 | 19243 | 338.47* | 387.07 | 343.70 | 91.5/78.2 | 3.2 |
| | HiFiCCL-Hifiasm | **3.29*** | **14525** | 441.76 | **459.30** | **399.82** | **91.9/83.0** | 3.2 |
| | HiFiCCL-Hifiasm (optional) | **3.27** | **14183*** | 451.49 | **464.17*** | **406.44*** | **91.9*/83.2*** | 3.2 |
| | HiFlye | 3.89 | 46896 | 855.19 | 462.76 | 361.91 | 92.4/81.0* | 2.4 |
| | HiFiCCL-HiFlye | 3.89 | 46988 | **832.09** | **492.60** | **388.44*** | **92.9***/80.9 | **2.1*** |
| | HiFiCCL-HiFlye (optional) | **3.88*** | **46726*** | **831.81*** | **493.24*** | **387.55** | **92.8**/81.0* | **2.3** |
| | LJA | 3.09 | 35718 | 120.08 | 148.37 | 138.94 | 78.7/69.4 | 11.1 |
| | HiFiCCL-LJA | 3.11 | 36083 | **91.84** | **153.38** | **145.13** | **80.5/70.8** | **9.5** |
| | HiFiCCL-LJA (optional) | 3.11 | 36090 | **90.60*** | **153.66*** | **145.54*** | **80.7*/71.0*** | **9.3*** |
| HG002 (HiFi 11x) | Hifiasm | 4.74 | 21109 | 466.63* | 605.84 | 524.14 | 96.1/59.4 | 1.4 |
| | HiFiCCL-Hifiasm | **4.03** | **14295*** | 715.97 | **819.08*** | **688.19*** | 95.9/**71.0** | 1.7 |
| | HiFiCCL-Hifiasm (optional) | **4.07** | **14957** | 689.11 | **814.66** | **682.23** | 95.9/**71.2*** | 1.6 |
| | HiFlye | 4.17 | 54508 | 971.73* | 663.17 | 519.05 | 94.1/78.8 | 1.8 |
| | HiFiCCL-HiFlye | 4.18 | 54972 | 981.62 | **763.90*** | **568.12** | **94.2***/78.8 | 2.0 |

| | | | | | | | | |
|---|---|---|---|---|---|---|---|---|
| | HiFiCCL-HiFlye (optional) | 4.18 | 55006 | 978.71 | **760.06** | **573.44*** | 94.1/78.8 | 2.1 |
| | LJA | 4.09 | 57160 | 106.64 | 123.48* | 116.53* | 81.2/58.2* | 5.7 |
| | HiFiCCL-LJA | 4.09 | 57640 | **95.12** | 121.38 | 114.48 | **81.4**/57.2 | 5.7 |
| | HiFiCCL-LJA (optional) | 4.08 | 57592 | **94.10*** | 121.33 | 114.52 | **81.5***/57.3 | 5.7 |

**Table S9. SVs detection in HG002 GIAB Tier 1 (high-confidence regions).**

| Dataset | Assembler | regions | Precision | Recall | F1 |
|---|---|---|---|---|---|
| HG002 (5x) | HiFiCCL-Hifiasm | [50, 1000] | 0.7506 | 0.6495 | 0.6964 |
| | | [1000, 3000] | 0.9466 | 0.6181 | 0.7479 |
| | | [3000, 7000] | 0.9741 | 0.5947 | 0.7385 |
| | | [7000, +∞] | 0.9193 | 0.5643 | 0.6993 |
| | Hifiasm | [50, 1000] | 0.7517 | 0.6466 | 0.6952 |
| | | [1000, 3000] | 0.9384 | 0.5883 | 0.7232 |
| | | [3000, 7000] | 0.9813 | 0.5526 | 0.7070 |
| | | [7000, +∞] | 0.9016 | 0.5445 | 0.6790 |
| HG002 (8x) | HiFiCCL-HiFlye | [50, 1000] | 0.7440 | 0.8191 | 0.7797 |
| | | [1000, 3000] | 0.4849 | 0.7599 | 0.5920 |
| | | [3000, 7000] | 0.4065 | 0.7842 | 0.5354 |
| | | [7000, +∞] | 0.3148 | 0.6732 | 0.4290 |
| | HiFlye | [50, 1000] | 0.7278 | 0.8204 | 0.7713 |
| | | [1000, 3000] | 0.4892 | 0.7649 | 0.5967 |
| | | [3000, 7000] | 0.3948 | 0.7710 | 0.5222 |
| | | [7000, +∞] | 0.2690 | 0.6633 | 0.3828 |
| | HiFiCCL-LJA | [50, 1000] | 0.8492 | 0.6454 | 0.7334 |
| | | [1000, 3000] | 0.8833 | 0.6119 | 0.7229 |
| | | [3000, 7000] | 0.8686 | 0.6263 | 0.7278 |
| | | [7000, +∞] | 0.8382 | 0.5643 | 0.6745 |
| | LJA | [50, 1000] | 0.8471 | 0.6312 | 0.7234 |
| | | [1000, 3000] | 0.8750 | 0.5833 | 0.7000 |
| | | [3000, 7000] | 0.8937 | 0.5973 | 0.7160 |
| | | [7000, +∞] | 0.8484 | 0.5544 | 0.6706 |

**Table S10. Comparison of large SVs detection using different assembly results and read alignment on HG002 GIAB Tier 1 (high-confidence regions)**

| Dataset | Assembler | Precision | Recall | F1 |
|---|---|---|---|---|
| HG002 (5x) | reads | 0.8769 | 0.5643 | 0.6867 |
| | Hifiasm | 0.9016 | 0.5445 | 0.6790 |
| | HiFiCCL-Hifiasm | 0.9193 | 0.5643 | 0.6993 |
| | HiFlye | 0.2761 | 0.6534 | 0.3882 |
| | LJA | 0.8750 | 0.3465 | 0.4964 |
| | Verkko | 0.7906 | 0.3366 | 0.4722 |

**Table S11. SVs detection on HG002 CMRG.**

| Dataset | Assembler | regions | Precision | Recall | F1 |
|---|---|---|---|---|---|
| HG002 (5x) | HiFiCCL-Hifiasm | [50, 1000] | 0.8728 | 0.5988 | 0.7103 |
| | | [1000, 3000] | 1.0000 | 0.5454 | 0.7058 |
| | | [3000, 5000] | 1.0000 | 0.2500 | 0.4000 |
| | | [5000, +∞] | 1.0000 | 0.6000 | 0.7499 |
| | | [50, +∞] | 0.8880 | 0.5862 | 0.7062 |
| | Hifiasm | [50, 1000] | 0.8655 | 0.5988 | 0.7079 |
| | | [1000, 3000] | 0.9166 | 0.5000 | 0.6470 |
| | | [3000, 5000] | 1.0000 | 0.2500 | 0.4000 |
| | | [5000, +∞] | 0.7500 | 0.6000 | 0.6666 |
| | | [50, +∞] | 0.8676 | 0.5812 | 0.6961 |
| HG002 (8x) | HiFiCCL-HiFlye | [50, 1000] | 0.9127 | 0.7906 | 0.8473 |
| | | [1000, 3000] | 0.7894 | 0.6818 | 0.7317 |
| | | [3000, 5000] | 1.0000 | 0.5000 | 0.6666 |
| | | [5000, +∞] | 0.8000 | 0.8000 | 0.8000 |
| | | [50, +∞] | 0.8971 | 0.7733 | 0.8306 |
| | HiFlye | [50, 1000] | 0.9078 | 0.7441 | 0.8178 |
| | | [1000, 3000] | 0.7647 | 0.5909 | 0.6666 |
| | | [3000, 5000] | 0.6000 | 0.7500 | 0.6666 |
| | | [5000, +∞] | 0.8000 | 0.8000 | 0.8000 |
| | | [50, +∞] | 0.8809 | 0.7290 | 0.7978 |
| | HiFiCCL-LJA | [50, 1000] | 0.9270 | 0.5174 | 0.6641 |
| | | [1000, 3000] | 1.0000 | 0.5000 | 0.6666 |
| | | [3000, 5000] | 1.0000 | 0.5000 | 0.6666 |
| | | [5000, +∞] | 1.0000 | 0.8000 | 0.8888 |
| | | [50, +∞] | 0.9380 | 0.5221 | 0.6708 |
| | LJA | [50, 1000] | 0.8541 | 0.4767 | 0.6119 |
| | | [1000, 3000] | 1.0000 | 0.4090 | 0.5806 |
| | | [3000, 5000] | 1.0000 | 0.5000 | 0.6666 |
| | | [5000, +∞] | 1.0000 | 0.8000 | 0.8888 |
| | | [50, +∞] | 0.8738 | 0.4778 | 0.6178 |

**Table S12. Comparison of large SVs detection using different assemblies and read alignment on HG002 CMRG.**

| Dataset | Assembler | Precision | Recall | F1 |
|---|---|---|---|---|
| HG002 (5x) | reads | 0.7500 | 0.6000 | 0.6666 |
| | Hifiasm | 0.7500 | 0.6000 | 0.6666 |
| | HiFiCCL-Hifiasm | 1.0000 | 0.6000 | 0.7499 |

**Table S13. Statistics of human genome scaffolding.** The bold data indicates that the HiFiCCL metric's performance surpassed that of the base assembler.

| Dataset | Assembler | Size (Gb) | Contigs number | MA | N50 (Mb) | NG50 (Mb) | Gene completeness (busco) | |
|---|---|---|---|---|---|---|---|---|
| | | | | | | | Complete /Single(%) | Missing (%) |
| HG002 (HiFi 5x) | HiFiCCL-Hifiasm | **2.79** | 1275 | **12822** | **136.52** | **124.67** | **89.1/85.6** | **5.8** |
| | Hifiasm | 2.73 | 1034 | 13133 | 135.39 | 123.25 | 88.2/84.9 | 6.3 |
| HG002 (HiFi 8x) | HiFiCCL-HiFlye | 3.89 | 34374 | **10382** | 136.45 | 148.57 | **96.0**/86.2 | **1.4** |
| | HiFlye | 3.89 | 33623 | 11131 | 137.28 | 150.89 | 95.8/86.2 | 1.6 |
| | HiFiCCL-LJA | 3.12 | 13582 | **5766** | **127.79** | **127.79** | **89.1/80.3** | **7.3** |
| | LJA | 3.09 | 13254 | 6750 | 126.57 | 125.54 | 86.8/78.4 | 9.1 |
| NA19240 (HiFi 5x) | HiFiCCL-Hifiasm | **2.64** | 1172 | **15823** | **129.61** | **119.72** | **85.5/82.6** | **7.9** |
| | Hifiasm | 2.58 | 983 | 16030 | 125.42 | 115.40 | 84.5/81.8 | 8.4 |
| NA19240 (HiFi 8x) | HiFiCCL-HiFlye | **4.10** | 40994 | **13864** | 139.11 | **153.85** | 95.6/**83.9** | 2.1 |
| | HiFlye | 4.11 | 40599 | 13949 | 139.92 | 153.72 | 95.6/83.7 | 1.9 |
| | HiFiCCL-LJA | 2.87 | **9630** | **7933** | 127.07 | 118.21 | 85.8/**78.8** | 9.6 |
| | LJA | 2.89 | 9815 | 8208 | 127.31 | 119.08 | 85.9/78.7 | 9.5 |

**Table S14. Statistics of chromosome-level scaffolds.** The bold data indicates that the HiFiCCL metric's performance surpassed that of the base assembler.

| Dataset | Assembler | Size (Gb) | Contigs number | MA | N50 (Mb) | NG50 (Mb) | Gene completeness (busco) | |
|---|---|---|---|---|---|---|---|---|
| | | | | | | | Complete /Single(%) | Missing (%) |
| HG002 (HiFi 5x) | HiFiCCL-Hifiasm | **2.70** | 24 | **12258** | **136.52** | **124.67** | **89.0/85.7** | **5.8** |
| | Hifiasm | 2.65 | 24 | 12639 | 135.39 | 123.25 | 88.2/85.0 | 6.3 |
| HG002 (HiFi 8x) | HiFiCCL-HiFlye | 3.03 | 24 | **9625** | 148.57 | 148.57 | **95.7/92.0** | **1.6** |
| | HiFlye | 3.05 | 24 | 10389 | 150.89 | 150.89 | 95.6/91.8 | 1.7 |
| | HiFiCCL-LJA | 2.62 | 24 | **5521** | **136.61** | **127.79** | **89.0/85.7** | **7.5** |
| | LJA | 2.60 | 24 | 6472 | 135.91 | 125.54 | 86.8/83.5 | 9.1 |
| NA19240( HiFi 5x) | HiFiCCL-Hifiasm | **2.56** | 23 | **15294** | **129.61** | **119.72** | **85.4/82.6** | **8.0** |
| | Hifiasm | 2.51 | 23 | 15582 | 125.42 | 115.40 | 84.5/81.9 | 8.4 |
| NA19240 (HiFi 8x) | HiFiCCL-HiFlye | 3.07 | 23 | **13185** | **153.85** | **153.85** | 95.1/**91.2** | 2.3 |
| | HiFlye | 3.08 | 23 | 13229 | 153.72 | 153.72 | 95.1/91.1 | 2.1 |
| | HiFiCCL-LJA | 2.53 | 23 | **7731** | 129.42 | 118.21 | 85.8/**82.7** | 9.6 |
| | LJA | 2.54 | 23 | 7977 | 129.79 | 119.08 | 85.8/82.6 | 9.5 |

**Table S15. Statistics of human genome scaffolding across different assemblies.**

| Dataset | Assembler | Size (Gb) | Contigs number | MA | N50 (Mb) | NG50 (Mb) | Gene completeness (busco) | |
|---|---|---|---|---|---|---|---|---|
| | | | | | | | Complete /Single(%) | Missing (%) |
| HG002 (HiFi 5x) | HiFiCCL-Hifiasm | 2.79 | 1275 | 12822 | 136.52 | 124.67 | 89.1/85.6 | 5.8 |
| | Hifiasm | 2.73 | 1034 | 13133 | 135.39 | 123.25 | 88.2/84.9 | 6.3 |
| | HiFlye | 3.35 | 18315 | 17881 | 136.48 | 136.48 | 90.0/84.1 | 4.9 |
| | LJA | 1.76 | 2620 | 7436 | 97.90 | 38.56 | 49.6/47.3 | 42.7 |
| | Verkko | 2.02 | 8454 | 30176 | 96.15 | 52.01 | 49.2/45.9 | 42.7 |
| NA19240 (HiFi 5x) | HiFiCCL-Hifiasm | 2.64 | 1172 | 15823 | 129.61 | 119.72 | 85.5/82.6 | 7.9 |
| | Hifiasm | 2.58 | 983 | 16030 | 125.42 | 115.40 | 84.5/81.8 | 8.4 |
| | HiFlye | 3.07 | 14194 | 23669 | 136.83 | 136.83 | 85.4/79.8 | 8.2 |
| | LJA | 0.64 | 1126 | 3785 | 31.48 | - | 23.0/22.3 | 72.1 |
| | Verkko | 1.06 | 5050 | 16998 | 45.20 | - | 26.7/23.2 | 63.1 |

**Table S16. Statistics of chromosome-level scaffolds across different assemblies.**

| Dataset | Assembler | Size (Gb) | Contigs number | MA | N50 (Mb) | NG50 (Mb) | Gene completeness (busco) | |
|---|---|---|---|---|---|---|---|---|
| | | | | | | | Complete /Single(%) | Missing (%) |
| HG002 (HiFi 5x) | HiFiCCL-Hifiasm | 2.70 | 24 | 12258 | 136.52 | 124.67 | 89.0/85.7 | 5.8 |
| | Hifiasm | 2.65 | 24 | 12639 | 135.39 | 123.25 | 88.2/85.0 | 6.3 |
| | HiFlye | 2.91 | 24 | 17290 | 147.30 | 136.48 | 89.7/86.0 | 5.1 |
| | LJA | 1.67 | 22 | 7270 | 97.90 | 38.56 | 49.4/47.8 | 43.0 |
| | Verkko | 1.81 | 24 | 29883 | 99.34 | 52.01 | 49.1/47.3 | 42.8 |
| NA19240 (HiFi 5x) | HiFiCCL-Hifiasm | 2.56 | 23 | 15294 | 129.61 | 119.72 | 85.4/82.6 | 8.0 |
| | Hifiasm | 2.51 | 23 | 15582 | 125.42 | 115.40 | 84.5/81.9 | 8.4 |
| | HiFlye | 2.72 | 23 | 23212 | 138.03 | 136.83 | 85.3/81.9 | 8.2 |
| | LJA | 0.59 | 22 | 3624 | 35.10 | - | 22.5/22.2 | 72.6 |
| | Verkko | 0.93 | 23 | 16800 | 50.03 | - | 30.3/29.4 | 62.3 |
